# Spatiotemporal dynamics of membrane surface charge regulates cell polarity and migration

**DOI:** 10.1101/2022.05.19.492577

**Authors:** Tatsat Banerjee, Debojyoti Biswas, Dhiman Sankar Pal, Yuchuan Miao, Pablo A. Iglesias, Peter N. Devreotes

## Abstract

During cell migration and polarization, hundreds of signal transduction and cytoskeletal components self-organize to generate localized protrusions. Although biochemical and genetic analyses have delineated many specific interactions, how the activation and localization of so many different molecules are spatiotemporally orchestrated at the subcellular level has remained unclear. Here we show that the regulation of negative surface charge on the inner leaflet of the plasma membrane plays an integrative role in the molecular interactions. Surface charge, or zeta potential, is transiently lowered at new protrusions and within cortical waves of Ras/PI3K/TORC2/F-actin network activation. Rapid alterations of inner leaflet anionic phospholipids, such as PI(4,5)P2, PI(3,4)P2, phosphatidylserine, and phosphatidic acid, collectively contribute to the surface charge changes. Abruptly reducing the surface charge by recruiting positively charged optogenetic actuators was sufficient to trigger the entire biochemical network, initiate *de novo* protrusions, and abrogate pre-existing polarity. These effects were blocked by genetic or pharmacological inhibitions of key signaling components such as Akt and PI3K/TORC2. Conversely, increasing the negative surface deactivated the network and locally suppressed chemoattractant-induced protrusions or subverted EGF-induced ERK activation. Computational simulations involving excitable biochemical networks demonstrated that slight changes in feedback loops, induced by recruitment of the actuators, could lead to outsized effects on system activation. We propose that key signaling network components act on, and are in turn acted upon, by surface charge, closing feedback loops which bring about the global-scale molecular self-organization required for spontaneous protrusion formation, cell migration, and polarity establishment.

## INTRODUCTION

Cell migration and polarity involve large networks of interacting signal transduction and cytoskeletal components ^1–4^. Several important features of these networks are emerging. First, there is an exquisite spatiotemporal coordination of the molecular events: For example, Ras, PI3K, and Rac activation along with actin polymerization define growing protrusions at the front of the cell while PTEN, RhoA, and myosin II leave these regions and return as the protrusions are withdrawn. To the extent that it has been examined, this complementary organization is conserved during macropinocytosis, phagocytosis, cytokinesis, within traveling cortical waves in different cells throughout phylogeny, and in apical-basal polarity of epithelial cells ^5–7^. Second, consistent with the network concept, multiple genetic, pharmacological, and mechanical perturbations on different individual components often result in surprisingly similar phenotypic changes suggesting that the overall “setpoint” of the network is more important than any single constituent ^8–19^. Although much has been gained by studying detailed interactions between different molecules, a systems level explanation is needed to account for these emergent properties. It is possible that the features could be a result of a series of stepwise protein-protein interactions but it is more likely that the organization depends on a biophysical property of the plasma membrane ^20^. Such global biophysical organizers have been suggested ^1, 11, 20^, but properties that track closely with the observed dynamic spatial and temporal organization, or that allow small alterations to bring about dramatic shifts in the setpoint of the entire system causing outsized phenotypic effects, have not been identified.

One prospective candidate for a such a biophysical organizer is inner membrane surface charge or zeta potential. The cell maintains a negative charge on the inner leaflet of the plasma membrane, primarily by regulating the concentrations of multiple anionic phospholipids^21–24^. Of course, numerous studies have focused on specific protein interactions with individual lipids such as PIP3^25, 26^ and many cytoskeletal proteins were reported to bear PI(4,5)P2 binding motifs^27^. Interestingly, it has been reported that enzymatic *lowering* of specific anionic lipids such as PI(4,5)P2 or PI(3,4)P2 can lead to the network *activation* and alterations in cell migration^9, 28^. Relatively few studies have focused on the role of membrane surface charge itself ^21, 29^. These have proposed that the zeta potential, in the range of −20 to −50 mV^30–32^, acts in regulating local pH and soluble second messenger concentrations, altering ion channel conductance, and controlling the association of peripheral membrane proteins ^21–24, 33–36^. It was also shown that the during immune synapse formation and phagocytosis, the membrane surface charge changes ^22, 37–39^. However, it was not envisioned or assessed whether network activation and protrusion formation are associated with increases or decreases in surface charge or whether direct perturbations of surface charge can activate/deactivate the network.

To address these questions, and to understand the possible role of surface charge as a biophysical organizer, we examined whether the surface charge profile changes during cell migration and polarity establishment and designed actuators to alter rapidly the surface charge in a particular domain of the membrane. Here, we first showed that PI(4,5)P2 and PI(3,4)P2, phosphatidylserine and phosphatidic acid are dynamically *lowered* during the *activation* of Ras/PI3k/Akt/F-Actin network. Based on these initial observations, we used a generic surface charge sensor to assess whether surface potential tracked with the activation process. We observed that the surface potential is high in resting cells and is transiently *lowered* during network *activation* and protrusion formation. Altering surface potential by recruiting our novel positively or negatively charged optogenetic actuators to the membrane in a spatiotemporally controlled fashion, we establish that the level of surface potential in a local membrane domain is not a consequence but sufficient and necessary for activation of the entire signal transduction network and directing migration and polarity phenotypes.

## RESULTS

To analyze the organization of coordinated signaling and cytoskeletal activities, we started by examining protrusion formation in migrating *Dictyostelium* cells as well as cortical wave propagation in *Dictyostelium* and RAW 264.7 macrophage cells (Figures S1A-S1C). As previously noted ^3, 5, 26, 40^, events that facilitate protrusion formation such as Ras/PI3K activation and actin polymerization localize to the front regions of migrating cells (Figures S1A and S1D), whereas those that suppress protrusions such as PTEN and myosin II localize to back (Figures S1E). An analogous complementarity was observed in the ventral wave patterns: the facilitative and suppressive components dynamically separate into broad, propagating “front-” and “back”-state regions, respectively (Figures S1F-S1I) ^6, 41–43^. The waves have been shown to control protrusion dimensions and corresponding modes of cell migration, and thus serve as a useful surrogate for “front/back” polarity. To quantitate the extent of correspondence or complementarity between the distributions of a pair of different molecular components, we developed a conditional probability-based index or CP index (methods; Figures S2A and S2B) which was positive or negative, respectively, for co-localized versus complementary-localized pairs (Figures S2C and S2D). For most of the data, both normalized line scans and CP indexes are reported to provide quantitative measures of fractional changes in biosensor levels as well as the extent of complementarity.

### PI(4,5)P2, PI(3,4)P2, PS, and PA dynamically co-segregate to back-state regions of the membrane

To gain insight into the spatiotemporal coordination among the signaling and cytoskeletal components, we examined several anionic phospholipids in the inner surface of the plasma membrane using genetically encoded biosensors. The distribution of PH_PLCδ_ indicated that PI(4,5)P2 was depleted in the front-state regions of ventral waves and from protrusions in migrating *Dictyostelium* cells (Figures 1A and 1B; Video S1). Equivalent patterns of PI(4,5)P2 were observed in macrophage ventral waves (Figure 1C; Figures S3A; Video S1), as previously reported^44, 45^. The PI(3,4)P2 biosensor, PH_CynA_, displayed a similar pattern in the *Dictyostelium* ventral waves (Figure 1D; Figures S3B; Video S2), consistent with our previous reports of its depletion from the protrusions^28, 46^. Line scans showed around 40-70% less PI(4,5)P2 and PI(3,4)P2 in front versus back regions of the waves. The consistently high negative values of CP indices for PI(4,5)P2 and PI(3,4)P2 showed that the complementarity with respect to PIP3 was maintained during the dynamic patterning (Figures 1E and 1F).

**Figure 1.**
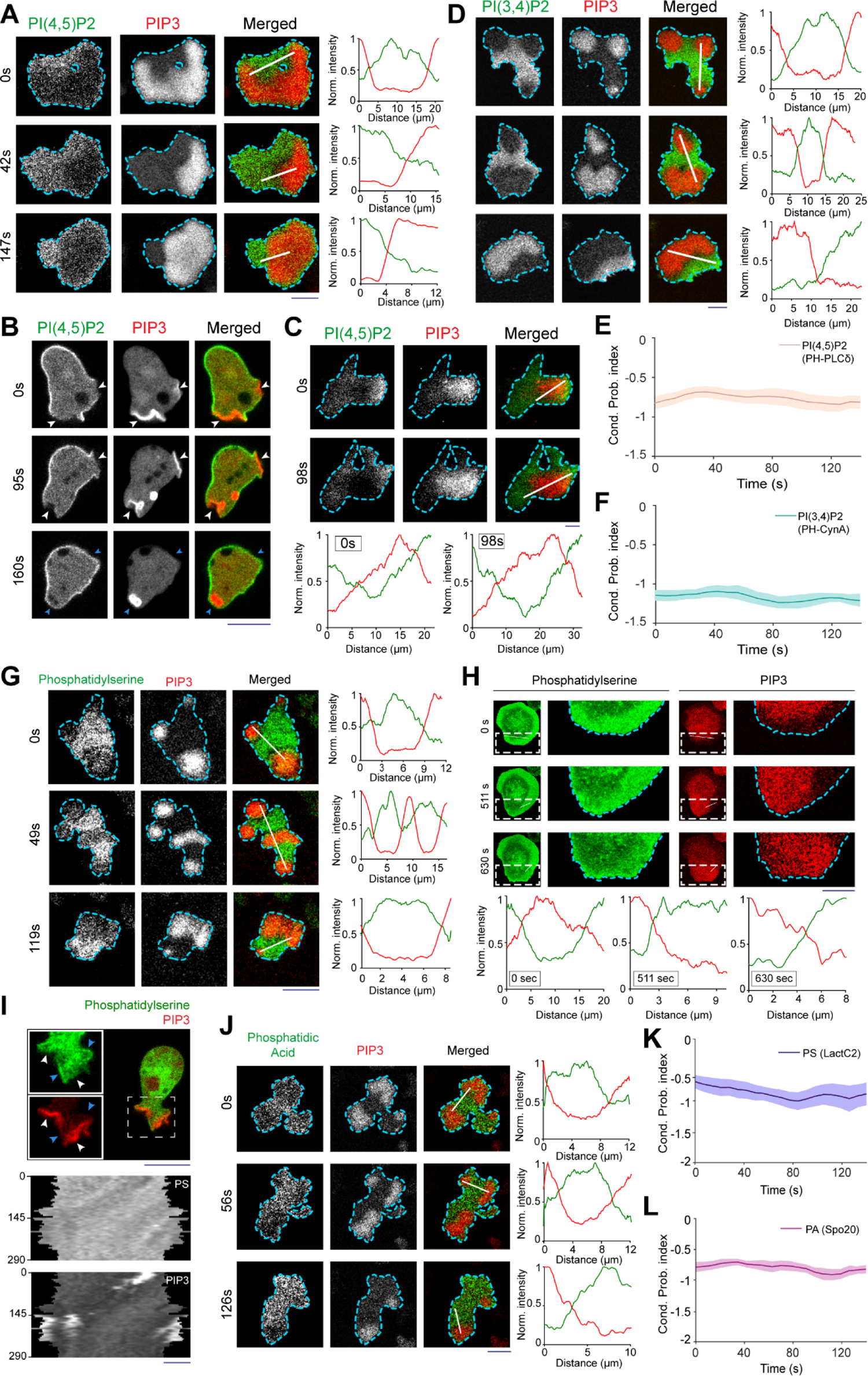
Multiple anionic phospholipids dynamically self-organizes to the back-state regions of the membrane. **(A and B)** Representative live-cell images of *Dictyostelium* cells co-expressing PI(4,5)P2 sensor PH_PLCδ_-GFP and PI(3,4,5)P3 sensor PH_crac_-mCherry, during ventral wave propagation (A) or migration (B). Note that PH_PLCδ_ dynamically localizes to the back-stare regions in ventral waves (A) and analogously moved away from protrusions in migrating cells (B). White arrows point to protrusions where PIP3 is enriched and PI(4,5)P2 sensor PH_PLCδ_ is depleted whereas blue arrows point to PH_PLCδ_ returning back to the membrane as protrusions were eventually retracted and membrane domain returned to its basal back-state (B). **(C)** Representative live-cell time-lapse images of RAW 264.7 macrophage co-expressing PI(4,5)P2 sensor GFP-PH_PLCδ_ and PI(3,4,5)P3 sensor PH_Akt_-mCherry demonstrating dynamic complimentary distribution in its ventral waves. **(D)** Representative live-cell time-lapse images of *Dictyostelium* cells co-expressing PI(3,4)P2 sensor PH_CynA_-KikGR and PIP3 sensor, demonstrating the spatiotemporal back localization of PI(3,4)P2 in its ventral waves. Corresponding linescans are displaying the intensity profile quantification in (A), (C), and (D). **(E and F)** Time series plot of CP index for PI(4,5)P2 (E) and PI(3,4)P2 (F). For all CP index time-plots in this paper, each of the n_c_ cells was analyzed for n_f_ =20 frames and all CP indices are calculated with respect to PIP3. PI(4,5)P2: n_c_=16 cells; PI(3,4)P2: n_c_=11 cells; data are mean ± SEM. **(G-I)** Dynamic back-state distribution of PS biosensor, GFP-LactC2, in ventral waves of *Dictyostelium* (G) and RAW 264.7 macrophages (H), and away from protrusions in migrating *Dictyostelium* cells (I). Protrusions and front-state regions of ventral waves are marked by PIP3 sensors. Bottom two panels of (I) show 360° membrane-kymographs around cell perimeter. **(J)** Complementary localization of PA sensor, GFP-Spo20, and PIP3 in *Dictyostelium* ventral waves. **(K and L)** Quantification of extent of back localization of PS and PA by time-series plot of CP index of LactC2 (K; n_f_=20 frames for each n_c_=15 cells) and Spo20 (L; n_f_=20 frames for each n_c_=16 cells); mean ± SEM. For all figures, scale bars are 10 μm.

Since these two signaling lipids segregated to the back-state regions of the membrane, we wondered whether other major anionic phospholipids, such as phosphatidylserine (PS) and phosphatidic acid (PA), exhibit asymmetric patterns. We studied the spatiotemporal distributions of PS, which accounts for ∼20% of inner leaflet lipids^47^, using its biosensor LactC2 ^47, 48^. Similar to PI(4,5)P2 and PI(3,4)P2, PS localized to the back-state regions of ventral waves of *Dictyostelium* (Figure 1G; Figures S3C; Video S3) and RAW 264.7 macrophages (Figure 1H; Figures S3D; Video S3) and was depleted from protrusions in migrating *Dictyostelium* cells (Figure 1I). Using the biosensor Spo20 ^49, 50^, we observed that PA preferentially distributed to the back-state regions of the ventral waves (Figure 1J; Figures S3E) and away from protrusions in migrating *Dictyostelium* cells (Figures S3F). Line scans showed around 30-50% less PS and PA in front versus back regions of the waves. The CP indices of PS and PA were also computed with respect to PIP3 (Figures 3K and 3L) and were found to be consistently negative, similar to PI(4,5)P2, PI(3,4)P2, and PTEN (Figures S3G). Taken together, these observations show that four major anionic lipids decrease in the front-state region of the membrane while PIP3 increases. While PIP3 carries the most negative charge, it is a minor constituent^47, 51–53^.

### Higher negative surface charge defines the inactive region or “back-state” regions of the cell cortex

Reasoning that, with respect to charge, the decreases in the four predominant anionic phospholipids would more than offset any increases in PIP3, we sought to measure spatiotemporal changes in membrane surface charge. We started with a generic membrane surface charge sensor, R(+8)-Pre, which had been quantitatively characterized *in vitro* (by liposome binding assays) to detect the combination of anionic lipids ^22, 30, 54^. Consistent with our observations of the individual lipids, we found that the back-state regions of cortical waves in *Dictyostelium* cells are defined by high negative surface charge whereas the front-state regions shun the charge sensor (Figure 2A; Video S4). Line scans demonstrated around 40-60% less charge sensor in front versus back regions of the waves (Figure 2A). Despite continuous propagation of front-region waves, as marked by PIP3, R(+8)-Pre faithfully maintained a preferential back-state localization (Figure 2B). Similarly, protrusions on migrating cells were depleted of charge sensor compared with the higher basal level on the membrane (Figure 2C; Video S4). An analogous spatiotemporal distribution of surface charge was also observed in the ventral waves of mammalian RAW 264.7 macrophage-like cells (Figures 2D and 2E; Video S5).

**Figure 2.**
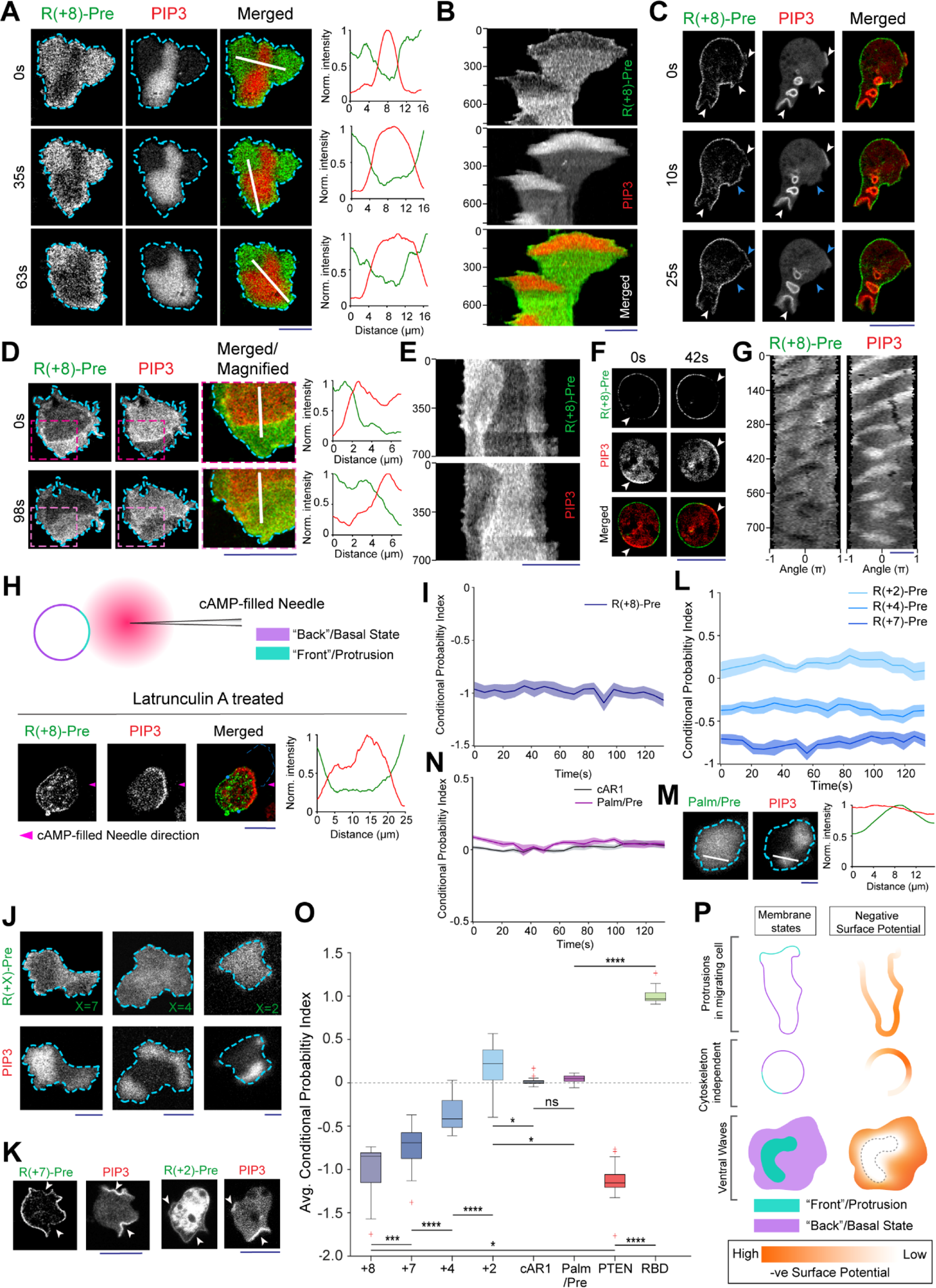
Back-state of the membrane maintains higher negative surface charge on the inner leaflet, compared to the front-state of the membrane. **(A)** Representative live-cell time-lapse imaging of a *Dictyostelium* cell co-expressing GFP-R(+8)-Pre and PIP3 biosensor, PH_Crac_-mCherry. **(B)** Representative line-kymograph of wave pattern shown in cell (A), showing the consistency of complementary localization of R(+8)-Pre and PIP3. In all kymographs, numbers on the left denote time in seconds, unless otherwise mentioned. **(C)** Live-cell time-lapse imaging of migrating *Dictyostelium* cell showing R(+8)-Pre is depleted in protrusions. White arrows: PIP3 enriched protrusion; Blue arrows: Retracted protrusions where PIP3 was depleted and R(+8)-Pre returned. **(D)** Ventral waves in RAW 264.7 macrophage cells co-expressing GFP-R(+8)-Pre and PIP3 biosensor, PH_Akt_-mCherry, displaying analogous complementary kinetics of the surface charge sensor with respect to PIP3. **(E)** Representative line-kymograph of wave pattern shown in cell (D), showing the consistency of the complementary localization. **(F and G)** Actin-polymerization inhibitor Latrunculin A treated *Dictyostelium* cells exhibiting the spatiotemporal surface charge gradient, complementary to PIP3. In (F), white arrows denote front-state consisting of bright patches PIP3 and depleted of surface charge sensor; (G) displayed the consistency of complementarity in a 360° membrane-kymograph around the periphery of the cell. **(H)** R(+8)-Pre was showing complementary localization with respect to front-state marker PIP3 during receptor activation mediated PIP3 production using chemotactic gradient stimulation. Magenta arrowhead indicates the direction of micropipette (filled with 1 μM cAMP) for gradient stimulation. Note that PIP3 was produced in the part of the membrane which is closer to the micropipette. Cells were pre-treated with Latrunculin A. **(I)** Time series plot of CP index of R(+8)-Pre, n_c_=30 cells; data are mean ± SEM. **(J)** Representative live-cell images of ventral waves in *Dictyostelium* cell membrane co-expressing PIP3 biosensor, PH_Crac_-mCherry, and GFP-R(+7)-Pre (left), GFP-R(+4)-Pre (middle), GFP-R(+2)-Pre (right). **(K)** Representative live-cell images showing R(+7)-Pre and R(+4)-Pre distribution, with respect to PIP3-rich protrusions, in migrating cells. **(L)** Time series plot of CP indices of mutated surface charge sensors R(+7)-, R(+4)-, R(+2)-Pre. +7: n_c_ =23, +4: n_c_ =20, +2: n_c_ =12 cells; data are mean ± SEM. **(M)** GFP-Palm/Pre is exhibiting uniform distribution in ventral waves, whereas PIP3 sensor PH_Crac_-mcherry is enriched in front-state regions. **(N)** Time series plot of CP indices of uniform membrane-markers cAR1 and Palm/Pre; cAR1: n_c_ =20, Palm/Pre: n_c_ =11, data are mean ± SEM. **(O)** Time averaged CP indices of surface charge sensors, uniform membrane marker controls, standard back protein PTEN (n_c_ =17), and front sensor RBD (n_c_ =15). To generate each datapoint, n_f_ =20 frames were averaged over these number of cells(n_c_). Box and whiskers are graphed using Tukey’s method. The p-values determined by Mann-Whitney-Wilcoxon test. **(P)** Schematic of negative surface potential distribution in front and back state of the cell during migration and cortical wave propagation.

Importantly, as previously reported for patterns of signal transduction events such as PIP3 accumulation and PTEN dissociation (Figures S1H) ^42, 43^, the dynamic distribution of surface charge was relatively independent of the actin cytoskeleton. In Latrunculin A-treated cells where periodic circulating waves were induced, the charge sensor continuously adjusted its localization towards the back-state during 12 minutes of observation (Figures 2F and 2G). It has been previously shown that cells exposed to a steady cAMP gradient, even when they are Latrunculin A treated, following a brief global response, display a crescent of front-region markers toward the high side of the gradient due to receptor-mediated activation of signaling events^55, 56^. As shown in Figure 2H and Figures S4A, when *Dictyostelium* cells were exposed to a chemotactic gradient, the charge sensor consistently moved away from the front region crescent and accumulated at the back. We quantified the complementarity between charge sensor and PIP3 in terms of CP index; in all these scenarios, the extent of complementary was found to be comparable to established back protein, PTEN (Figure 2I; Figures S2D).

These asymmetric distributions depended on charge and not on the specific amino acid sequences of the sensor. As the positively charged arginines of the charge sensor were sequentially replaced with neutral glutamines (Table S1), the distribution became increasingly uniform over the cortex (Figures 2J-2L; Figures S4B-S4F). The dynamics of R(+2)-Pre sensor (Figures 2J and 2K; Figures S4D and S4F) resembled the uniform distribution of surface receptor cAR1 (Figures S4G and S4H), the weakly charged lipidated sequence of LYN (Figures S4I), and the uncharged lipidated C-terminal tail of HRas (Figure 2M; Figures S4J), as quantified in terms of CP indices (Figures 2L, 2N, and 2O). In contrast, two distinct polybasic sequences, PTEN_1-18_-CAAX and RacG_CT_ (Table S1), carrying +6 and +7 charges, respectively, displayed a complementarity to front sensor in ventral waves and protrusions, which was roughly correlated with their net amount of charge (Figures S5A-S5G). All these data collectively suggest that, under various physiological scenarios, such as in protrusions in migrating cells, in activated crescents of Latrunculin A treated cells, and in activated regions of propagating ventral waves, the front-state of the membrane maintains a lower negative surface charge, compared to the back-state (Figure 2P).

### Anionic lipids contribute to the surface charge in an integrative manner

As multiple anionic lipids decreased during the activated phase, we tested whether any single phosphoinositide fully accounted for the asymmetric distribution of the surface charge sensor. When we rapidly depleted PI(4,5)P2 by recruiting Inp54p to the membrane, PH_PLCδ_, moved to the cytosol (Figure 3A; Figures S6A), as expected. However, R(+8)-Pre remained membrane bound (Figures 3B and 3C). As previously published^9, 57^, upon PI(4,5)P2 depletion, the cells switched migration mode and displayed increased velocity (Figure 3D; Figures S6B and S6C). Correspondingly, the front- and back-state regions expanded and contracted, respectively.

Nevertheless, R(+8)-Pre still faithfully localized to the back-state of the cortex (Figure 3E). Similarly, in *Dictyostelium Dd5p4*^-^ cells, where PI(3,4)P2 levels are low, front regions are expanded, and migration is also altered ^28^, R(+8)-Pre remained on the membrane (Figures S6D and S6E) and maintained its characteristic distribution to the remaining back-state regions of the cell (Figure 3F). To be certain that membrane retention of surface charge sensor following recruitment of Inp54p was not due to its binding to the product PI4P, we recruited pseudojanin by a similar chemically induced dimerization system. This particular chimera of INPP5E and sac1 was reported to be sufficient to simultaneously convert PI(4,5)P2 to PI4P and PI4P to PI^58^. As expected, we found that upon recruitment of pseudojanin from cytosol to membrane, PI(4,5)P2 sensor PH_PLCδ_ moved to cytosol (Figure 3G and Video S6) and so did PI4P-specific sensor PHOsh2X2 (Figure 3H and Video S6). However, almost all of the R(+8)-Pre remained on the membrane (Figures 3I and 3J; Video S7). Together, these results demonstrate that the dynamic regulation of the membrane surface charge depends on PI(4,5)P2 and PI(3,4)P2, but there are additional key contributors, such as PS and PA, for example.

We sought a possible mechanism for the local decreases of PS on activated regions of inner membrane since it is not readily modified by kinase/phosphatases like the phosphoinositides. Reversible flipping of PS in non-apoptotic conditions has been previously reported in several physiological contexts such as cytosolic calcium release and immunological stimulation^59, 60^. However, despite multiple speculations over the years^22, 47^, whether rapid and dynamic externalization can regulate local decreases in PS pertinent to signaling and cytoskeleton activation has not been shown. We performed a transient Annexin-V binding assay (see methods for details) to label outer leaflet PS during protrusion formation. Remarkably, we found that, on the outer leaflet, PS strongly localizes to the protrusion or front-state regions(Figure 3K), closely opposing its inner leaflet localization (Figure 1G-1I). We confirmed that in quiescent cells where protrusions were absent, Annexin V did not bind to outer leaflet (Figures S6F). Pearson’s correlation coefficient analysis demonstrates that extent of co-localization of front-state or protrusion marker LimE and Annexin V is very similar to the extent of co-localization of two standard front-state makers such as PH_Crac_ and RBD (Figure 3L) and significantly different than standard complementary pair such as PH_Crac_ and PTEN (Figure 3L). Obviously, more studies will be required, but our data here suggest that inner leaflet PS may transiently flips to outer leaflet in the membrane domains where the biochemical network is activated or where protrusions form.

As explained earlier, in the inner leaflet of the membrane, PI(3,4,5)P3 is present at negligible amounts compared to other anionic phospholipids^47, 51–53^ and theoretically, it can perform its well-known signaling functions without significant contribution to surface charge^52^. Nevertheless, we directly tested the contribution of PIP3 to the membrane surface charge profile using genetic and pharmacological perturbations. First, we treated the cells with LY294002, a potent inhibitor of PI3K^61–64^, which resulted in a loss of PIP3 accumulation in the protrusions (Figure 3M). Yet, the typical depletion of the surface charge sensor from the protrusions was largely unchanged (Figure 3M). Second, we used *Dictyostelium PI3K 1^-^/2^-^* cells where PIP3 production is severely impaired^62, 65^. Observing the surface charge sensor profile in those cells with respect to different front-state events such as F-actin polymerization (Figure 3N) and Ras-activation (Figure 3O), we again found that negative surface charge is consistently lowered during signaling and cytoskeletal network activation, irrespective of PI(3,4,5) levels. These results demonstrate that PIP3 does not significantly contribute to the membrane surface charge.

**Figure 3.**
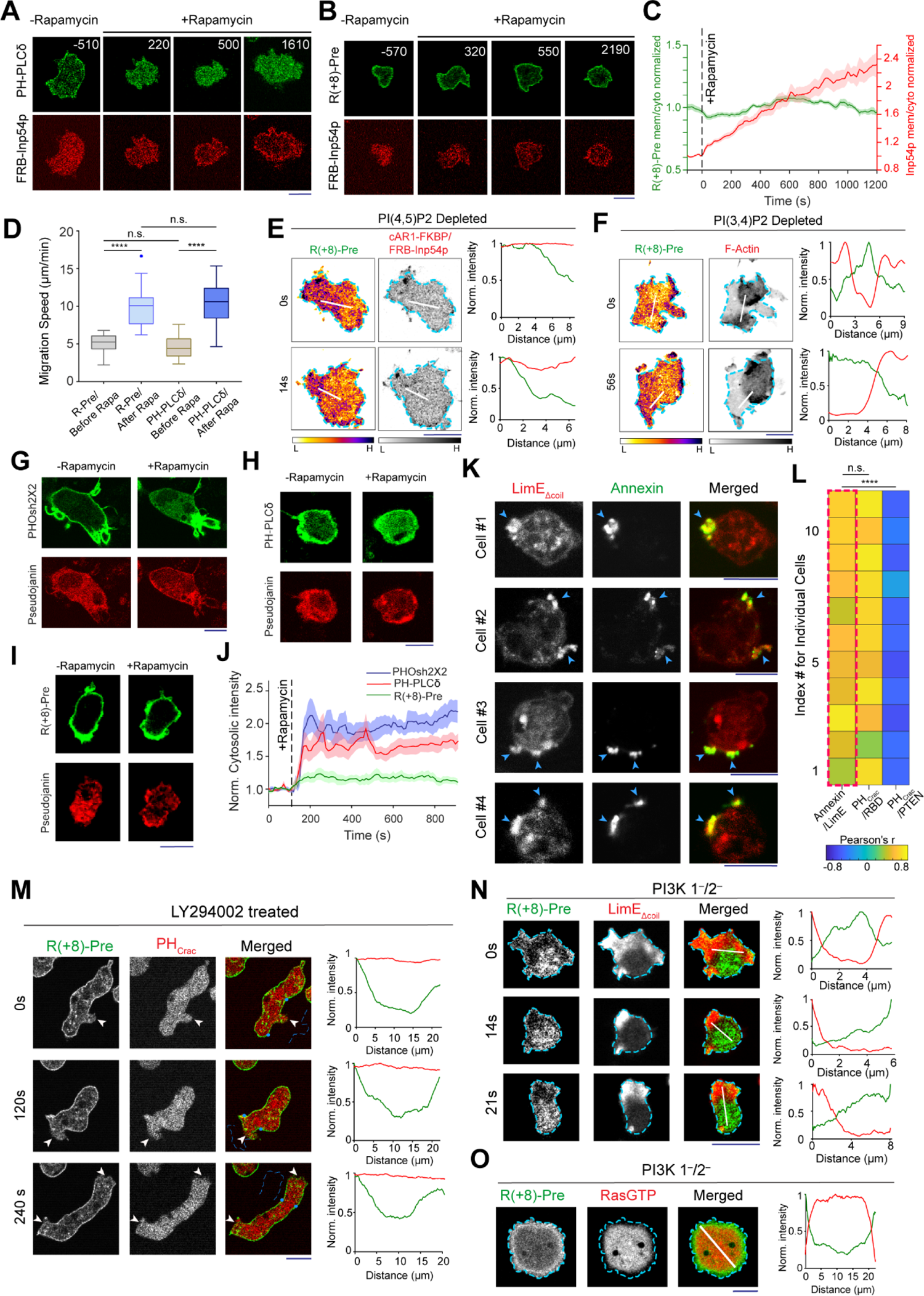
Dynamics of inner membrane surface charge sensor R(+8)-Pre in different anionic phospholipid depleted cells. **(A and B)** Representative live-cell time-lapse images showing the dynamics of membrane localization of PH_PLCδ_-GFP (A) or GFP-R(+8)-Pre (B), before and after PI(4,5)P2 depletion in *Dictyostelium* cells, co-expressing the chemically induced dimerization system i.e. cAR1-FKBP and mCherry-FRB-Inp54p. Numbers indicate time in seconds; rapamycin added at 0s. **(C)** Time course of membrane-to-cytosol ratio of R(+8)-Pre and Inp54p upon rapamycin addition (indicated by black dashed vertical line), demonstrating R(+8)-Pre did not dissociate from membrane upon PI(4,5)P2 depletion; n_c_=20 cells; mean ± SEM. **(D)** Migration speed of cells, co-expressing CID system for PI(4,5)P2 depletion, along with either GFP-R(+8)-Pre or PH_PLCδ_-GFP, before and after rapamycin addition; n_c_=32 cells tracked for n_f_=60 frames (7s/frame) for each case; p-values by Mann–Whitney–Wilcoxon test. **(E)** Spatiotemporal back localization of R(+8)-Pre in PI(4,5)P2 depleted cells, shown in “fire invert” colormap of Fiji/ImageJ; Inp54p recruited to uniform membrane-anchor cAR1 is symmetric in ventral waves. **(F)** Complementary localization of R(+8)-Pre with respect to front marker LimE (biosensor for newly-polymerized F-Actin) in ventral waves of Dd5p4^-^ *Dictyostelium* cells. **(G-I)** Representative live-cell images of RAW 264.7 macrophages showing the membrane localization profile of PI4P biosensor PHOsh2X2-GFP(G), PI(4,5)P2 biosensor PH_PLCδ_-GFP (H), or GFP-R(+8)-Pre (I), before and after recruiting Pseudojanin to membrane anchor Lyn-FRB. **(J)** Time course of normalized cytosolic intensity of PHOsh2X2, PH_PLCδ_, and R(+8)-Pre, upon rapamycin addition; time of addition is indicated by black dashed vertical line; n_c_=16 cells for PHOsh2X2, n_c_=10 cells for PH_PLCδ_, n_c_=12 cells for R(+8)-Pre; mean ± SEM. **(K)** Representative examples of protrusion forming *Dictyostelium* cells expressing LimE_Δcoil_-GFP, whose outer leaflet of membrane was allowed to transiently bind with Annexin V. (L) Heatmap of Pearson correlation coefficient between Annexin V and LimE_Δcoil_ (leftmost column, shown with dashed red line). Pearson correlation coefficient between (PH_Crac_ and RBD) and (PH_Crac_ and PTEN) are shown to demonstrate standard Co- and counter-localization profiles. All correlation coefficients were calculated along cell membrane; n_c_=11 cells in each case; p-values by Mann-Whitney-Wilcoxon test. **(M)** Representative live-cell time-lapse images of migrating *Dictyostelium* cell co-expressing GFP-R(+8)-Pre and PIP3 sensor PH_Crac_-mCherry which was treated with PI3K inhibitor LY294002. White arrows: protrusions which were depleted of PIP3 (due to LY294002 treatment), yet maintained surface charge gradient. **(N and O)** Representative live-cell time-lapse images of ventral waves in *PI3K 1^—^/2^—^ Dictyostelium* cells co-expressing GFP-R(+8)-Pre and front-state marker LimE (N) or RBD (O), demonstrating surface charge sensor is dynamically distributing to the back-state regions of the membrane.

### Decrease in net membrane surface potential is sufficient to trigger protrusions and abrogate the pre-existing polarity

The coordinated regulation of generic surface charge suggests that it is a key integrator of signal transduction and cytoskeletal networks. Previously, the spatiotemporal behavior of these networks has been modeled as an excitable network consisting of positive and delayed negative feedback loops ^9, 57, 66–69^. Independent studies suggest that the delayed negative feedback loop required for excitability includes the action of substrates of AKTs and possibly other refractory state molecules^57, 70^. The positive feedback was implemented as a mutually inhibitory interaction between front and back states, consisting of Ras/Rap and PI(4,5)P2 or PI(3,4)P2, respectively ^9, 28, 57^. Our data suggest that, while decreases in specific lipids can be effective activators, mutually inhibitory positive feedback loop may be controlled by overall surface charge (Figure 4A). In this scenario, the activated state would decrease negative charge while decreasing negative charge would lead to activation. Having demonstrated the former, we next sought to assess whether directly perturbing the negative surface charge on the membrane could alter the strength of the loop and thereby affect protrusion formation, polarity organization, and migratory behavior.

**Figure 4.**
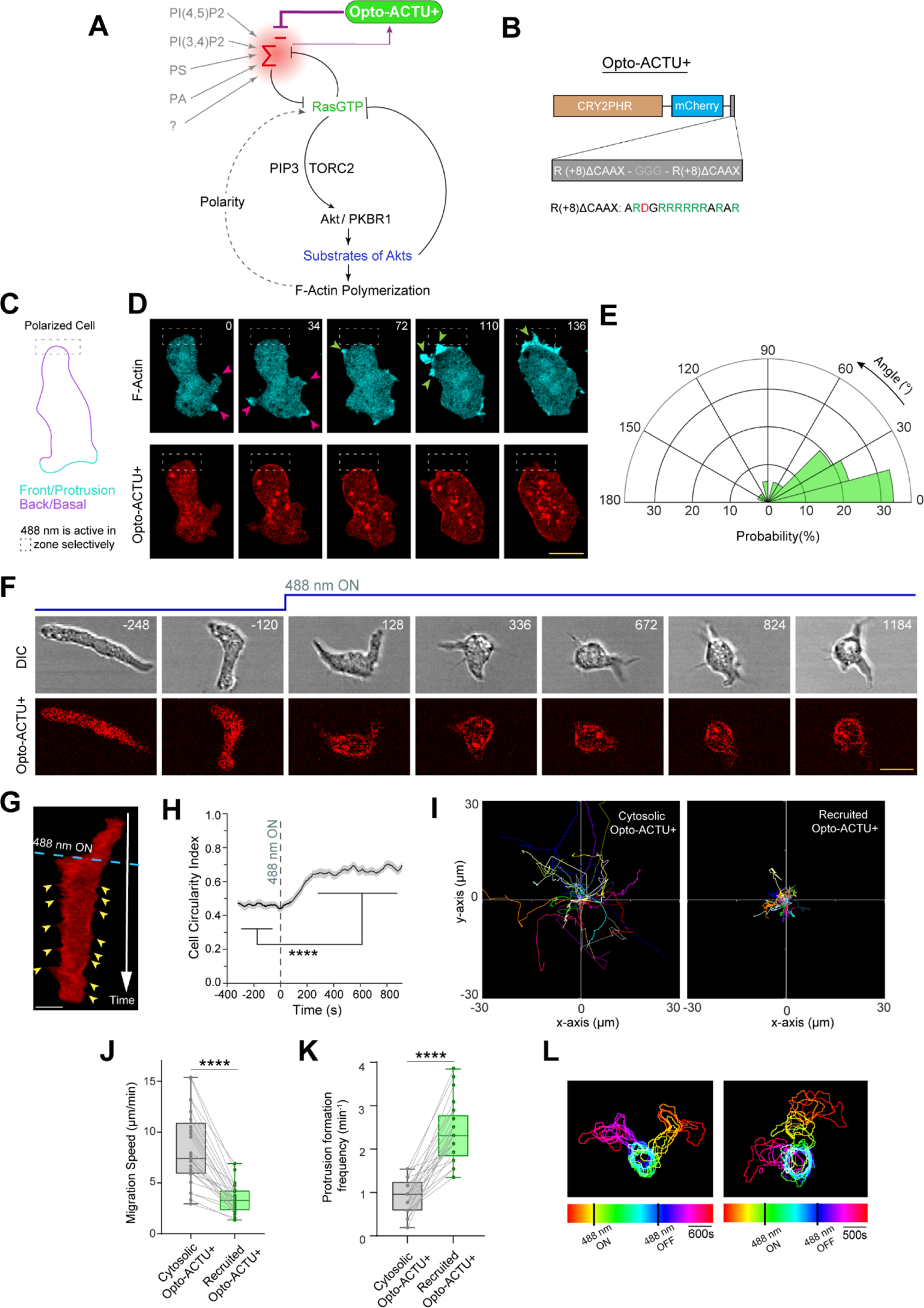
Lowering negative surface charge on the membrane can trigger *de novo* generation of protrusions and can abolish the pre-existing polarity. (A) Scheme for lowering negative surface charge by the recruitment of positively-charged optogenetic-actuator, Opto-ACTU+, in context of biochemical excitable network topology. Σ^—^: back-state defined by overall negative surface charge. Opto-ACTU+ interfere into the topology by getting associated with Σ^—^ but in turn it provides a negative feedback to Σ^—^. **(B)** Design of Opto-ACTU+ with net charge +16. Positively-charged amino acids in green, negatively-charged amino acids in red. **(C)** Setup of selective optical recruitment at the back of polarized *Dictyostelium* cells. **(D)** Representative time-lapse images of selective *de novo* protrusion formation from the area of recruitment in *Dictyostelium* cells co-expressing Opto-ACTU+, cAR1-CIBN, and LimE-Halo. Dashed rectangle shows the area where 488 nm laser was selectively illuminated for recruitment. Magenta and green arrows show existing and newly induced protrusions, respectively. Time in seconds. **(E)** Polar histogram of angle of protrusion formation with respect to recruitment area, demonstrating the higher probability of protrusion formation near the area of recruitment; n_c_=23 cells, n_p_=36 protrusions. **(F and G)** Time-lapse snapshots (F) and time-stack (G) demonstrating cell morphology and migration mode changes in a polarized *Dictyostelium* cell, upon recruitment of Opto-ACTU+. Cells were co-expressing Opto-ACTU+ and cAR1-CIBN. Numbers are time in seconds. 488 nm laser switched ON *globally* at t=0s. Yellow arrowheads: newly generated protrusions. Note that Opto-ACTU+ is depleted in the protrusions. **(H-K)** Quantification of cell morphology and migration mode changes in terms of cell circularity index (H), cell tracks (I), migration speed (J), and new protrusion formation frequency (K) upon Opto-ACTU+ recruitment (n=25 cells). Polarity loss shown as mean ± SEM over time (H). In (I-K), for either before or after recruitment, each cell tracked for n_f_=40 frames (8 sec/frame was the image acquisition frequency). Tracks were reset to the same origin in (I). For pairwise comparison, tracks are color-coded in (I) and data from same cell are connected by gray lines in (J) and (K). **(L)** Two representative examples of temporally color-coded cell outlines showing cell morphology and migratory mode before 488nm was turned on, during 488nm kept on, and after 488nm was switched off. All p-values by Mann-Whitney-Wilcoxon test.

To this end, we developed two non-specific biophysical optogenetic actuators designed to acutely increase or decrease the negative surface potential profile. Unlike previously reported actuators that were developed based on specific components such as GEFs, GAPs, RGS, or kinases/phosphatases ^71–75^, these unique actuators, consisting of a short chain of charged amino acids, are not directed at any particular biochemical reaction.

The first actuator, designated Opto-ACTU+, had positive charge of +16 (Figure 4B). We anticipated that when recruited from cytosol to the membrane in a spatially restricted fashion (Figure 4C; Figures S7A), it would reduce the negative surface charge and thereby increase positive feedback (Figure 4A). As a result, the threshold of the excitable networks would be lower, the probability of spontaneous firing would increase, and consequently, the cell would generate more protrusions *de novo*. Indeed, when Opto-ACTU+ was recruited to a quiescent back region of a polarized *Dictyostelium* cell, new protrusions started forming nearby (Figure 4D; Figures S7B; Video S8), in a remarkably similar fashion as previously obtained by specific biochemical perturbations such as optical activation of Rac1, Cdc42, or the GPCR opsins ^76–78^. With recruitment of uncharged Opto-CTRL new protrusions rarely appeared near the irradiation area (Figures S7C and S7D; Video S8), indicating that the protrusion formation is due to the negative surface charge reduction and not light irradiation or cryptochrome recruitment. As the angular histograms demonstrate, the probability of new protrusion generation was highest in the vicinity of the Opto-ACTU+ recruitment (Figure 4E) whereas Opto-CTRL recruitment did not induce a consistent bias (Figures S7E).

We anticipated that when Opto-ACTU+ is globally recruited from cytosol to membrane in polarized cells, more protrusions would be generated, and polarity would be disrupted. Indeed, within few minutes of global recruitment of Opto-ACTU+, cells began to rapidly extend protrusions all along the cortex (Figures 4F and 4G; Video S9), including from domains of erstwhile back-states. Consequently, polarity was abrogated (Figure 4H) and migration was impaired (Figures 4I and 4J). The number of new protrusions increased around 2.5 fold (Figure 4K) which is consistent with amount of surface charge reduction (Supplementary Note 1; methods). The entire series of events was reversible: When the 488 nm laser was turned off, the Opto-ACTU+ returned to the cytosol, cells repolarized and resumed polarized migration (Figure 4L; Video S10). No consistent changes in polarity, migration, or protrusion formations were observed upon the recruitment of uncharged control Opto-CTRL (Figures S8A and S8B; Video S9), as quantified in terms of cell circularity index (Figures S8C), speed (Figures S8D and S8E), and protrusion formation frequency (Figures S8F).

Careful observation revealed that in addition to more protrusion formation induced by the recruitment of the Opto-ACTU+, a series of events was occurring: As new protrusions/front-state regions eastablished, recruited Opto-ACTU+ was quickly moving away from these regions (Figures 4F and 4G; Video S9). In other words, once recruited, Opto-ACTU+ accumulated in a back-state region, it triggered a protrusion there, and subsequently translocated away, presumably due to the decreased negative surface charge in these regions. It gradually accumulated inside the newly formed back region and the entire cycle was repeated multiple times (Figures S8G).

### The effects of surface charge alteration on migration and polarity are mediated by signaling and cytoskeleton network components

To investigate the molecular mechanism through which membrane surface charge acts, we employed a series of genetically encoded biosensors, different pharmacological and genetic perturbations, in conjunction with our optogenetic actuators (Figure 5A). First, when Opto-ACTU+ was recruited, it accumulated to the existing back-state regions of the cortex and actin polymerization was eventually initiated there (Figure 5B). We observed that Opto-ACTU+ accumulation in a specific domain of the membrane consistently preceded the F-actin polymerization (Figure 5B; Figures S9A and S9B). Moreover, after formation of a newly polymerized actin-based protrusion, Opto-ACTU+ quickly moved away (Figures S9A) and accumulated to the newly forming back which, again, gradually started F-actin polymerization that led to protrusion (Figures 5B and Figures S9B). Next, we monitored the activation and localization kinetics of different signaling molecules upon surface charge perturbation, which are upstream regulators of F-actin based membrane protrusions. We started with PTEN, an established back-protein. The first 18 amino acids of PTEN, which were reported to be essential for membrane binding^79, 80^, when lipidated are sufficient to localize to the back-state region of the cortex, presumably due to its +6 charge (Figures S5B, S5D, and S5F; Table S1). We speculated that when Opto-ACTU+ would be recruited and accumulated to a domain of the membrane, surface charge would be lowered there, and as a result, owing to the charge-sensitive domain, PTEN would dissociate from that particular domain. Indeed, localization of Opto-ACTU+ to a spatially confined region within the back-state region of the membrane consistently caused a dissociation of PTEN from that region (Figure 5C and Figures S9C). Such spatiotemporally regulated loss of PTEN contributed to a dynamic accumulation of PI(3,4,5)P3 inside those transient Opto-ACTU+ enriched domains (Figures S9D and S9E). Since Ras superfamily GTPases are considered to activate PI3K (resulting in PI(3,4,5)P3 production) and TORC2 (Figure 5A), we tested whether Ras was activated upon surface charge lowering. Using an RBD biosensor that indicates the GTP-bound state of Ras, we found that Ras was dynamically activated in the surface charge lowered regions throughout the time course of the experiment (Figure 5D and Figures S9F).

**Figure 5.**
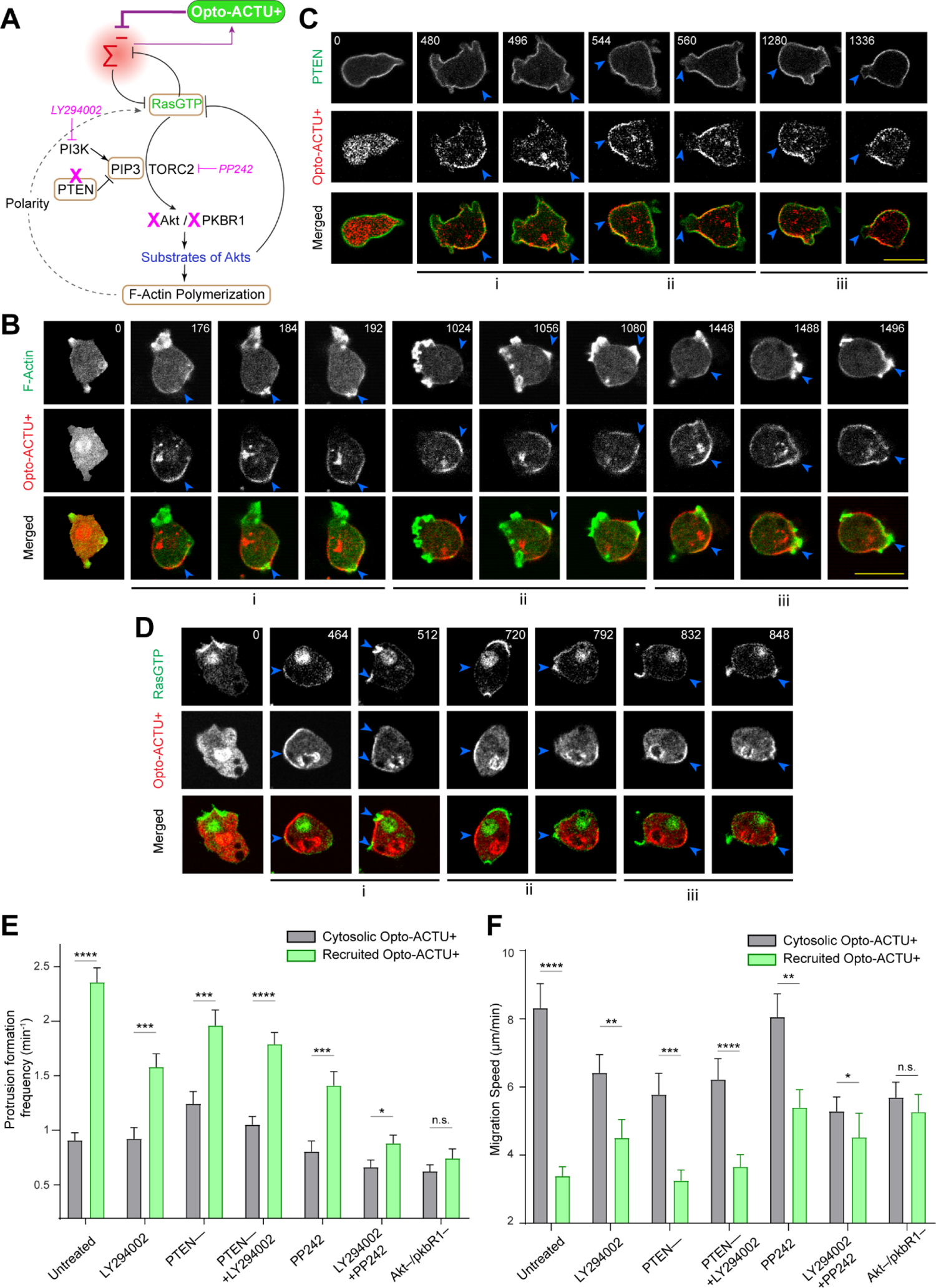
The phenotypic changes induced by Opto-ACTU+ recruitment are mediated by the Ras-PI3K/Akt/TORC2/F-actin network. **(A)** Scheme showing nodes of signaling and cytoskeletal network that were monitored and/or perturbed, in conjunction with Opto-ACTU+ recruitment. Tan-colored rectangles denote the molecules whose dynamics were recorded. Magenta blocked arrows denote pharmacological inhibitions and magenta cross-marks denote genetic knockouts. **(B)** Time-lapse live cell images of *Dictyostelium* cells co-expressing Opto-ACTU+, cAR1-CIBN, and LimE-GFP (biosensor for newly polymerized F-actin), where recruitment was started at t=0s (first time point). Throughout this figure, numbers on images show time in seconds. The “i”, “ii”, “iii” are showing three representative actin polymerization cases. For each case, three events are shown: first, Opto-ACTU+ accumulated inside a domain of the membrane; second, F-actin polymerization was initiated there; and finally, when that domain fully turned into front state, Opto-ACTU+ moved away from that domain. Throughout this figure, blue arrowheads in are showing the domains of interest, i.e. where Opto-ACTU+ was first accumulated. **(C)** Time-lapse live cell images of *Dictyostelium* cells co-expressing Opto-ACTU+, cAR1-CIBN, and PTEN-GFP, where recruitment was started at t=0s (first time point). The “i”, “ii”, “iii” are showing three representative cases of PTEN dissociation from the membrane. For each case, two events are shown: first, the accumulation of recruited Opto-ACTU+ inside a domain of the membrane and second, when that resulted in the dissociation of PTEN from that particular domain of the membrane which also caused Opto-ACTU+ to move away from there. **(D)** Time-lapse live cell images of *Dictyostelium* cells co-expressing Opto-ACTU+, cAR1-CIBN, and RBD-GFP (biosensor for activated Ras), where recruitment was started at t=0s (first time point). The “i”, “ii”, “iii” are showing three representative cases of PTEN dissociation from the membrane. For each case, two events are shown: first, the accumulation of recruited Opto-ACTU+ inside a domain of the membrane and second, when that resulted in the activation of Ras inside that particular domain of the membrane which also caused Opto-ACTU+ to move away from there. **(E and F)** Quantification of phenotypic changes upon Opto-ACTU+ recruitment in terms of new protrusion formation frequency (E) and migration speed (F), in presence of different pharmacological inhibitors or genetic knockouts. Untreated: n_c_= 28 cells; LY294002: n_c_= 28 cells; PP242: n_c_= 22 cells; PTEN –: n_c_= 22 cells; PTEN –+ LY294002: n_c_= 24 cells; LY294002+PP242: n_c_= 27 cells; Akt^—^/PKBR1^—^ double knockout: n_c_= 21 cells. For each case, each of the n_c_ cells were tracked for n_f_=40 frames (8 sec/frame was imaging frequency) and time averages were taken. The mean ± SEM are shown. For pairwise comparison and more detailed data, please see Figures S10. The p-values by Mann-Whitney-Wilcoxon test.

Next, we sought to modify different nodes of Ras/PI3K/TORC2/Akt/F-Actin network using different genetic knockouts and drug treatments to interfere with the Opto-ACTU+-mediated increased protrusion formation and consequent loss of polarity and migration speed (Figure 5A). First, to investigate the role of PTEN in the signaling network activation, we performed Opto-ACTU+ recruitment experiment in *pten*– *Dictyostelium* cells which have elevated PIP3 levels and exhibits impaired migration profile and higher protrusion numbers^79, 81, 82^. We observed that recruitment of Opto-ACTU+ in developed *pten*– cells was sufficient to induce even more frequent protrusion formation (Figure 5E; Figures S10A) which resulted severely impaired migration profile (Figure 5F; Figures S10B and S10C). We next tested the effects of reduced PIP3 level by pre-treating *Dictyostelium* cells with the PI3K inhibitor LY294002^61–64^. PI3K inhibition only slightly inhibited the Opto-ACTU+ recruitment driven migration phenotypic changes: cells still made more protrusions (Figure 5E; Figures S10D) and thus migrated less (Figure 5F; Figures S10E). To confirm that the membrane surface charge perturbation mediated signaling and cytoskeleton network activation are not merely dependent on PTEN or PIP3-centric mechanisms, we treated the *pten*– cells with LY294002 and then performed Opto-ACTU+ recruitment. We found that, even when both PTEN and PI3K activity are impaired, cells consistently displayed increased activity in terms of protrusion formation (Figure 5E; Figures S10F), migrated slowly (Figure 5F; Figures S10G and S10H), and the polarity was abrogated (Figures S10I and Video S11), upon Opto-ACTU+ recruitment. Similarly, standalone inhibition of TORC2 by PP242^9, 83^ could only slightly hinder the Opto-ACTU+ recruitment driven migration phenotype changes (Figures 5E and 5F; Figures S10J and S10K), compared to untreated condition (Figures 5E and 5F; Figures 4J and 4K). On the other hand, inhibition of both PI3K and TORC2 by the cocktail of LY294002 and PP242 almost completely blocked the change in migration speed and protrusion formation (Figures 5E and 5F; Figures S10L and S10M). It is well established that the activities of PI3K and TORC2 converge to activate AKT and AKT-related kinase, PKBR1, that together activate numerous regulators of cytoskeleton^9, 62, 63^. Since AKT and PKBR1 act in a redundant fashion in different physiological scenarios^62^, we used *Akt ^-^/PKBR1^-^ Dictyostelium* cells to study the effects of surface potential perturbation. We observed that in these cells, Opto-ACTU+ recruitment did not induce any significant changes either in terms of protrusion formation rate (Figure 5E; Figures S10N) or migration mode (Figure 5F; Figures S10O). Together, these findings suggest that surface charge-mediated changes in migration or protrusion formation acts via the collective action of Ras/PI3K/TORC2/AKT/F-Actin network.

### Simulations of actuators dynamics in excitable network capture the phenotypic changes resulting from global and local lowering of surface charge

To simulate these unique polarity breaking and protrusion formation phenomena, we incorporated separate actuator dynamics into a model of the excitable network that includes polarity ^84^ (methods; Figures 6A-6C). Prior to perturbing the system with the actuators, polarity biased the excitable network, resulting in persistently localized firings which underlie the protrusions (Figure 6D). Following simulated recruitment, Opto-ACTU+ lowered the threshold and caused abrupt increases in overall activity along the whole perimeter. With the development of each front-state region, Opto-ACTU+ quickly distributed to new back-state regions where it reduced the local threshold, resulting in triggering of new protrusions (Figure 6D). As in experiments, this cycle repeated multiple times. The reversibility of this actuation process was also recreated in the simulations by allowing the actuators to dissociate from the membrane (Figures S11A and S11B). We also simulated the selective protrusion formation by confining the recruitment of our actuator within a back region and observed increased activity there which resulted a switch in polarity and, as the actuator was consistently cleared out from the activated area (Figure 6F). Together, these results suggest that, during polarized cell migration, higher negative surface charge at the back of the cell lead to an increased threshold which prevents protrusion formation there.

**Figure 6.**
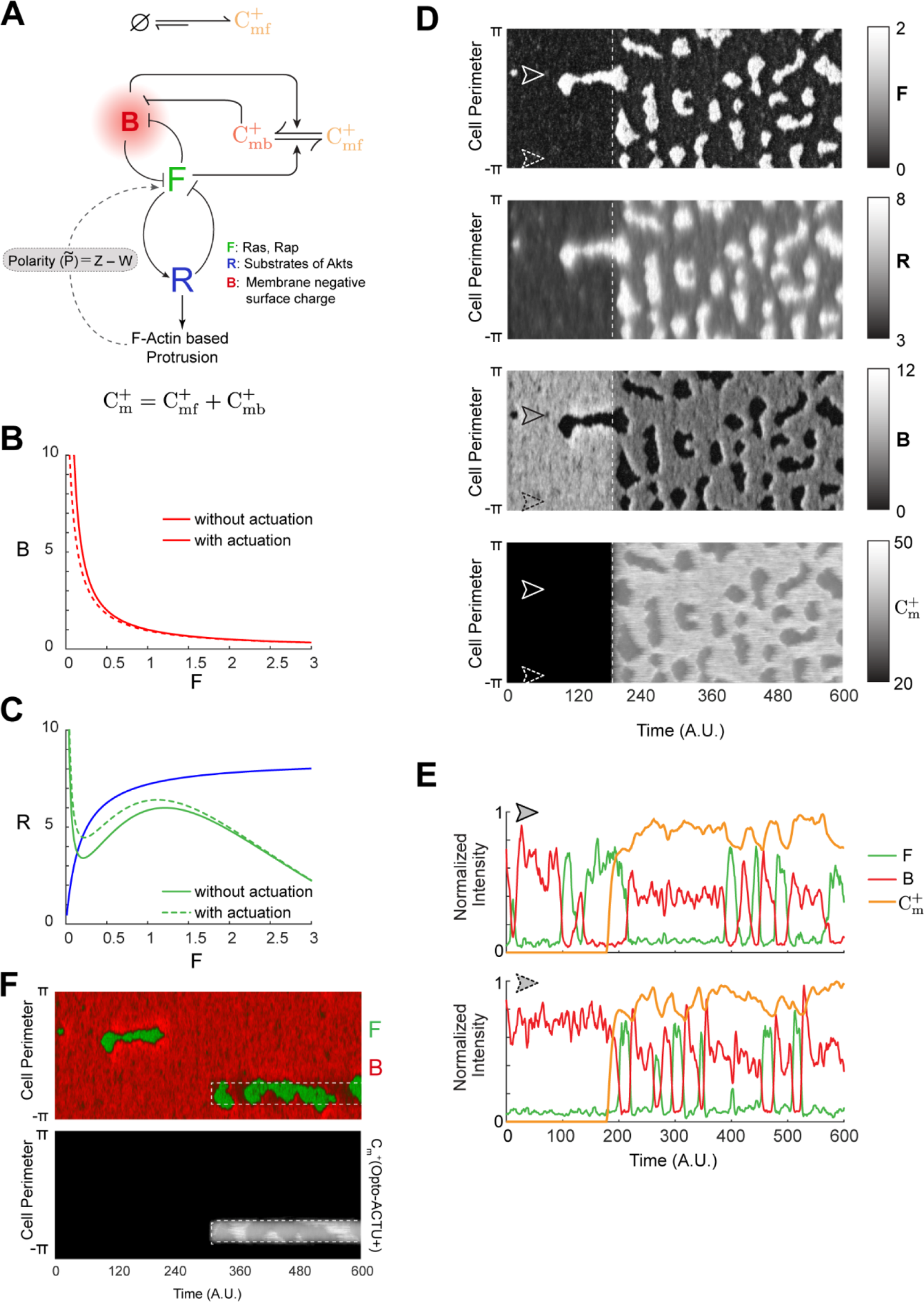
*In silico* lowering of surface charge recreates the polarity breaking and demonstrates increased activity over the membrane. **(A)** Schematic showing coupled system of excitable network, polarity module (involving Z and W), and Opto-ACTU+ (*C^+^_m_*) system. The excitable network involves membrane states F (front), B (back, defined by overall surface charge of inner membrane), and R (refractory). The polarity module comprises of local activator Z and delayed globally diffusing inhibitor W. The Opto-ACTU+ system constitutes of fast diffusing state *C_mf_*^+^ and almost stationary membrane bound state *C_mb_*^+^. The total charge actuator, *C_m_*^+^ on the membrane is the summation of both the states. **(B)** Plot of B vs F with and without Opto-ACTU+. **(C)** F and R nullclines with and without Opto-ACTU+ (under the steady-state assumption for B). **(D)** The simulated kymographs of F (first), R (second), B (third) and *C_m_*^+^ (fourth) in response to global recruitment. The instant of recruitment is shown by the white dashed line. **(E)** Line scans at two locations (denoted by arrows) on the simulated kymographs showing the temporal profiles of F (green), B (red) and *C_m_*^+^ (orange). **(F)** Simulated kymographs of membrane states in response to the selective recruitment of Opto-ACTU+ (*C_m_*^+^). Merged view of F (green) and B (red) are shown in the top panel; profile of Opto-ACTU+ (*C_m_*^+^) was shown in the bottom panel (in grayscale); the location of *in silico* selective recruitment is denoted by the white dashed box. **(G)** Schematic showing the cationic concentration (*C_m_*^+^ /Ca^2+^) elevation system coupled with the excitable network and the polarity module. **(H)** The simulated kymographs in response to global increase in *in silico* calcium concentration. Merged view of F (green) and B (red) are shown in the top panel and the profile of *C_m_*^+^ /Ca^2+^ was shown in the bottom panel.

### Spatially confined elevation of surface charge is sufficient suppress protrusions locally

Since reducing negative surface charge was able to activate the signaling network, we asked whether direct elevation of negative surface charge could deactivate the network and thereby limit protrusions (Figure 7A). To test this idea, we designed a second optogenetic peptide, Opto-ACTU-, with net negative charge of −14 that is expected to quickly increase the negative surface charge on the membrane upon recruitment (Figure 7B). We chose mammalian macrophage-like RAW 264.7 cells that are generally unpolarized and quiescent but can be globally activated by C5a receptor agonist so that they make protrusions all along the membrane. We first recruited Opto-ACTU- to a region on the membrane to increase negative surface charge locally and then globally stimulated the cell with C5a receptor agonist (Figure 7C). Protrusions formed along the membrane except in the vicinity of Opto-ACTU-recruitment region (Figure 7D; Figures S12A; Video S12). This induced a polarity in the cell and the cell migrated away (Figure 7D; Figures S12A; Video S12). Thus, selective surface charge manipulation can produce migration patterns that resemble those previously elicited by direct perturbations of the signaling and cytoskeletal network components^8, 17, 85^. Compared to Opto-ACTU-(Figure 7E), local recruitment of uncharged Opto-CTRL could not suppress protrusions upon global agonist stimulation (Figures S12B; Video S12) and the probability of protrusion formation remained uniform (Figures S12C). In simulations, we first elevated the threshold locally mimicking Opto-ACTU-recruitment and superimposed a global reduction of threshold representing the addition of receptor agonist (Figures S12D). We found increased activity except where the local suppression was enforced (Figure 7F, Figures S12E).

**Figure 7.**
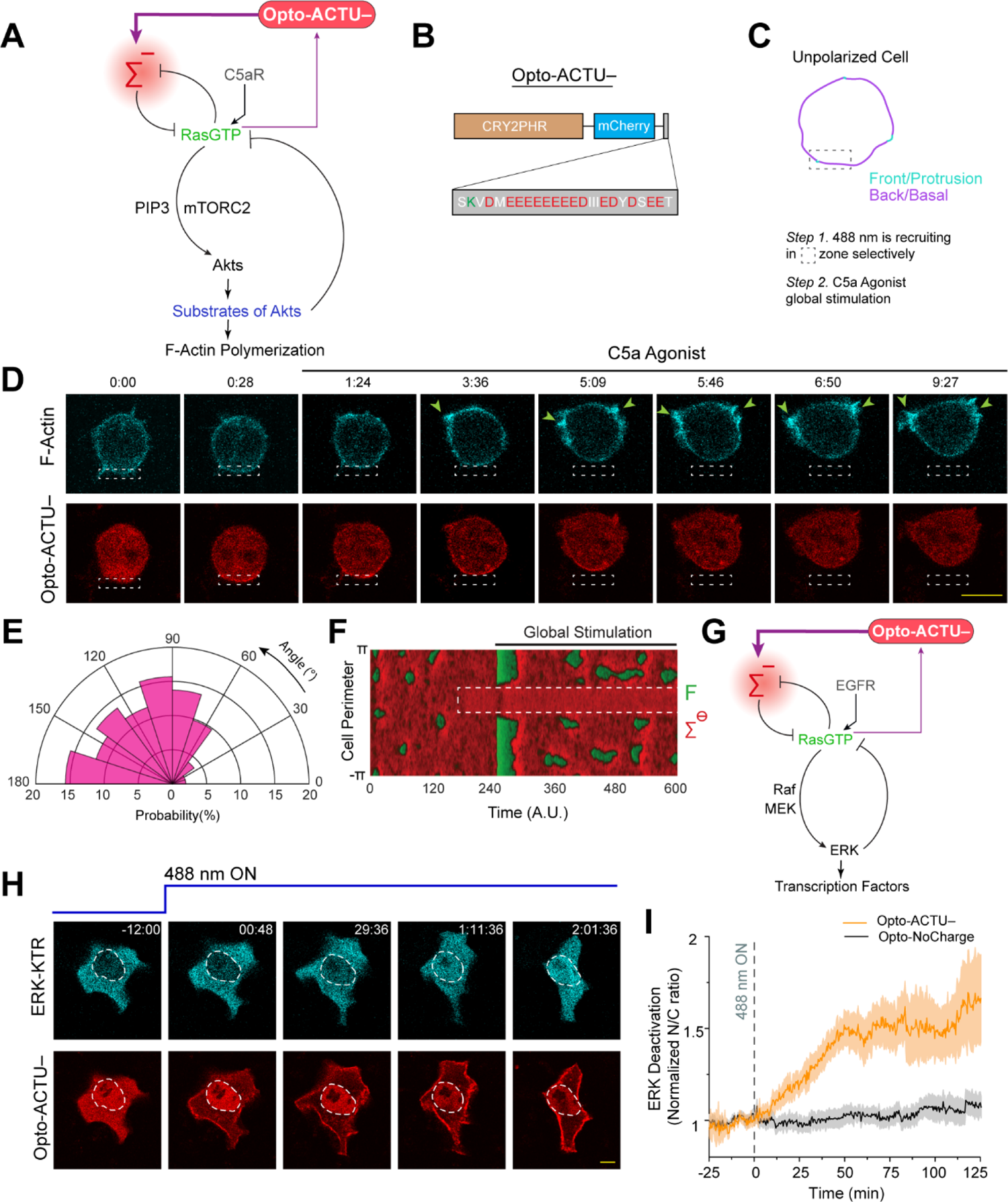
Increase in negative surface potential in the membrane suppresses protrusions and, separately, deactivates the EGF induced ERK activity. **(A)** Scheme for elevation of negative surface charge on membrane by the recruitment of negatively-charged optogenetic actuator, Opto-ACTU–, in context of biochemical excitable network topology with receptor input. **(B)** Design of Opto-ACTU-with net charge −14. Positively-charged amino acids in green, negatively-charged amino acids in red. **(C-E)** Experimental setup of selective Opto-ACTU– recruitment, followed by uniform C5a stimulation, in unpolarized RAW 264.7 macrophages (C); representative time-lapse images demonstrating cell migration driven by selective protrusion suppression in the site where Opto-ACTU- was locally recruited and protrusion formation in other areas of cortex upon uniform C5a stimulation (D); polar histogram indicating higher probability of protrusion formation away from recruitment area. Time in min:sec format (D); Green arrows: F-actin-rich protrusions marked by Lifeact (d); n_c_=12 cells, n_p_=55 protrusions (E). Cells were co-expressing Opto-ACTU–, CIBN-CAAX, and Lifeact-mVenus. **(F)** Simulated kymograph of membrane states in response to the *in silico* recruitment of Opto-ACTU– (C ^-^), followed by global stimulation. Front or F-state is in green, back or Σ^-^-state is in red. **(G)** Scheme for elevation of negative surface charge in context of excitable network-mediated ERK regulation, along with receptor input module. **(H)** Representative time-lapse images of epithelial MCF10A cell displaying ERKKTR translocation from cytosol to nucleus demonstrating ERK deactivation, upon Opto-ACTU– global recruitment to membrane; cells were pre-treated with and maintained in a saturating dose of EGF throughout the experiment; time in hr:min:sec format (H). **(I)** Quantification of ERK deactivation in terms of ERKKTR nucleus/cytosol ratio; n=12 cells for each case (Opto-ACTU– and Opto-CTRL), mean ± SEM; 488 nm was turned on first at t=0 min.

### Global increase in membrane surface charge is sufficient to subvert EGF induced ERK activation in epithelial cells

We next asked whether increasing negative surface potential on the membrane can override the EGF receptor mediated activation of ERK since the pathway involves activation of Ras (Figure 7G). Mammary epithelial cells, MCF10A, were first activated with a saturating dose of EGF, which was confirmed by the predominantly cytosolic distribution of ERK activation sensor ERK-KTR (Figure 7H first time point; Figures S12F first time point, Video S13). When we raised the negative surface charge of the membrane by globally recruiting Opto-ACTU-(Figure 7G), a substantial fraction of ERK-KTR became nuclear, indicating a deactivation of ERK (Figure 7H; Video S13), whereas control Opto-CTRL recruitment caused no visible effect (Figures S12F; Video S13). Quantitation of multiple cells showed that, although there were fluctuations of ERK activity (Figures S12G and S12H), Opto-ACTU-recruitment provoked ∼50% increase in nucleus/cytosol ratio of KTR, compared to uncharged control (Figure 7I).

## DISSCUSSION

Our study demonstrates that negative surface charge on the inner leaflet of the plasma membrane spatiotemporally coordinates the components of the Ras/PI3K/TORC2/F-actin network that control cell migration and polarity. We show that, under physiological conditions, the levels of phosphoinositides PI(4,5)P2 and PI(3,4)P2 as well phosphatidylserine and phosphatidic acid on the inner membrane decrease during network activation at membrane protrusions and front-state regions on ventral surfaces of cells. These changes collectively account for a major reduction in membrane surface charge or zeta potential which serves to integrate the signal transduction events in those activated regions. Altering membrane surface charge is necessary and sufficient for network activation and, not a merely a consequence of it, since recruiting a positively charged actuator to the membrane activates the network whereas recruiting a negatively charged actuator prevents the chemoattractant or growth factor stimulated activation. Our results suggest that inner membrane surface charge is a key biophysical parameter that appears within the feedback loops that determines the set-point of these networks.

Our data suggest that the fluid-mosaic model of the membrane is more complex than it is originally envisioned. We find that the inner leaflet of the plasma membrane is constantly remodeling in a highly coordinated manner. Large patches of multiple anionic lipids and peripheral membrane proteins co-segregate into defined “phase” domains which propagate dynamically. In the case of the phosphoinositides and many of the peripheral membrane proteins, the propagation is not due to lateral translocation but rather a sequential transient biochemical modification of headgroups of the lipids and shuttling of the proteins^68, 86, 87^. Many of the enzymes that modify the headgroups of phosphoinositides have been identified and for a handful of cases, their specific activation cascade is known. For example, PIP3 levels on the membrane increases when PI3K is activated by Ras^88, 89^ as well as when PTEN level decreases^81, 90^.

However, in-depth future studies will be needed to determine which specific kinases and phosphatases are activated or inhibited in sequence to control the levels of PI(4,5)P2, PI(3,4)P2 or PA in membrane. Additionally, as our data suggest, the regulation of PS is possibly regulated by dynamic externalization and hence we anticipate that several flippases, floppases, and scramblases whose activation tracks closely with Ras/PI3K/TORC2/F-Actin network activation and which co-segregate in front or back-state phase, will be identified in future.

The optogenetic actuators we developed allowed us to alter inner membrane surface charge in a generic fashion, inside an intact live cell, in a spatiotemporally restricted fashion. Thus, this induced, biophysical change, i.e. a reduction in zeta potential, sets in motion a series of biochemical events that were previously shown to be involved in migration, protrusion formation, and polarity. Despite its implications, such an effect of surface charge perturbation on different signaling and cytoskeletal events was hitherto largely unexplored, primarily due to dearth of available tools to directly perturb inner membrane surface charge. Traditional approaches with isolated membranes ^29, 31, 37, 91–94^ or giant unilamellar vesicle (GUV)/liposome or charged surfactant usually cannot accurately mimic the physiological environment of the cell due to incorrect lipid compositions, lack of counterions, and/or compromised inner leaflet integrity. Other recent methods^22, 24, 47, 52^, such as phosphoinositide degradation/synthesis by chemically induced dimerization or pharmacological perturbations to increase cytosolic cation concentrations or cellular ATP depletion do not provide any spatial control. Moreover, these approaches do not decouple the surface charge from other cell physiological changes.

Optogenetics do provide spatial and temporal control inside intact live cells, but most have been engineered to directly change the biochemical activity and/or localization of one particular network component^71, 72, 75^.all of those were engineered to directly change the biochemical activity and/or localization of one particular network component. As far as we know, the synthetic optogenetic systems we have developed here are the first to assess the effect of standalone surface charge perturbation *in situ*. Since these novel actuators can work orthogonally to external cues such as chemoattractant gradients or external electric field, we envision that these can potentially help in developing more in-depth molecular architecture of receptor mediated biochemical network activation. We anticipate that in different physiological scenarios such as immune synapse formation and phagocytosis where surface charge remodeling has been reported (and others that likely will be found in the future), our tools will enable unique perturbation strategies that can reveal intricate coordination pattern.

Our study establishes negative surface potential as an organizer of signal transduction and cytoskeletal events that control cell migration and polarity. We suggest the term “action surface potentials” to describe the traveling membrane domains of transiently decreased negative surface charge. We observed that high negative surface charge corresponds to a “resting” or back-state of the membrane while the regions of decreasing surface charge are transitioning to an active/front-state which leads to protrusions. We envision that, as surface charge decreases, molecules that regulate the signaling activities respond differentially to charge. Further studies are needed to determine which other crucial network components may be directly regulated by charge. We speculate that these changes in turn initiate downstream events which mediate further loss of multiple anionic lipids in the front regions and possibly other events, additionally decreasing the membrane surface charge. This arrangement constitutes a positive feedback loop that enables small fluctuations such as a small initial drop in zeta potential to expand into propagating waves and can have outsized phenotypic effects. This situation would be analogous to the ability of transmembrane potential to regulate key ion channels which in turn regulates transmembrane potential during action potential propagation.

The “action surface potential” hypothesis can explain a series of heretofore puzzling observations: First, as we discussed, in most signal transduction networks a vast number of activities undergo a highly coordinated stereotypical transient response. This extraordinary degree of coordination within an extensive series of stepwise interactions could be facilitated if the regulation of key components depended on an organizer such as the surface charge. Second, cells expend significant energy to sustain an asymmetric distribution of anionic lipids on inner leaflet of the membrane, which has little apparent structural advantage. Of course, PIP2 serves as a substrate for PLC and PI3K and PS as a signal for apoptosis. However, we suggest that another, perhaps larger, role of this asymmetric distribution of charged lipids is to help in setting up a basal state for triggering of action surface potentials which involve changes in multiple lipids acting in a common direction. Third, the excitable nature of the action surface potentials enables global control over cytoskeletal activities and underlies oscillations for frequency control of gene expression. The lateral propagation of the waves along the cortex in more or less circular patterns provide form and dimension to protrusions. Recent reports highlight that travelling waves of cytoskeletal and signaling activities mediate a diverse range of cell physiological processes in different kinds of cells and organisms^13, 68, 69, 86, 95–100^. Our results imply that action surface potential likely contribute to spatiotemporally orchestrating these events.

## Acknowledgments

We thank S. Grinstein (Hospital for Sick Children/University of Toronto) for helpful discussions. We are grateful to M. Kozlov (Tel Aviv University) for helping us with resident charge calculation. We thank all members of Devreotes and Iglesias laboratories as well as members of D. Robinson and M. Iijima laboratories (Johns Hopkins University School of Medicine) for valuable suggestions. We are grateful to Y. Long and Y. Deng (Devreotes lab) for helping with some experiments. We would like to thank N. Gautam (Washington University School of Medicine in St. Louis), O. D. Weiner (UCSF), and R. R. Kay (MRC LMB) for providing cells. We thank G. Du (McGovern Medical School, UTHealth) and A. Müller-Taubenberger (LMU Munich) for sharing plasmids. We thank Addgene and dictyBase for providing all the plasmids and resources. This work was supported by NIH grant R35 GM118177 (to P.N.D.), DARPA HR0011-16-C-0139 (to P.A.I. and P.N.D.), AFOSR MURI FA95501610052 (to P.N.D.), as well as NIH grant S10 OD016374 (to S. Kuo of the JHU Microscope Facility).

## Author contributions

T.B. and P.N.D. conceptualized the overall study. T.B. designed and performed all *Dictyostelium* experiments. D.S.P. introduced the mammalian culture and T.B. and D.S.P. together designed and carried out the mammalian experiments. Y.M. provided resources and contributed to the experiments. D.B. and P.A.I. developed the software to compute the conditional probability index and performed localization analysis. T.B. quantified and analyzed other results, with inputs from other authors. D.B. and P.A.I. developed the computational models and D.B. conducted all the simulations. T.B., P.N.D., D.S.P., D.B., and P.A.I. wrote the manuscript. Second authors are listed alphabetically. P.N.D. and P.A.I. supervised the study.

---

**Figure S1.**
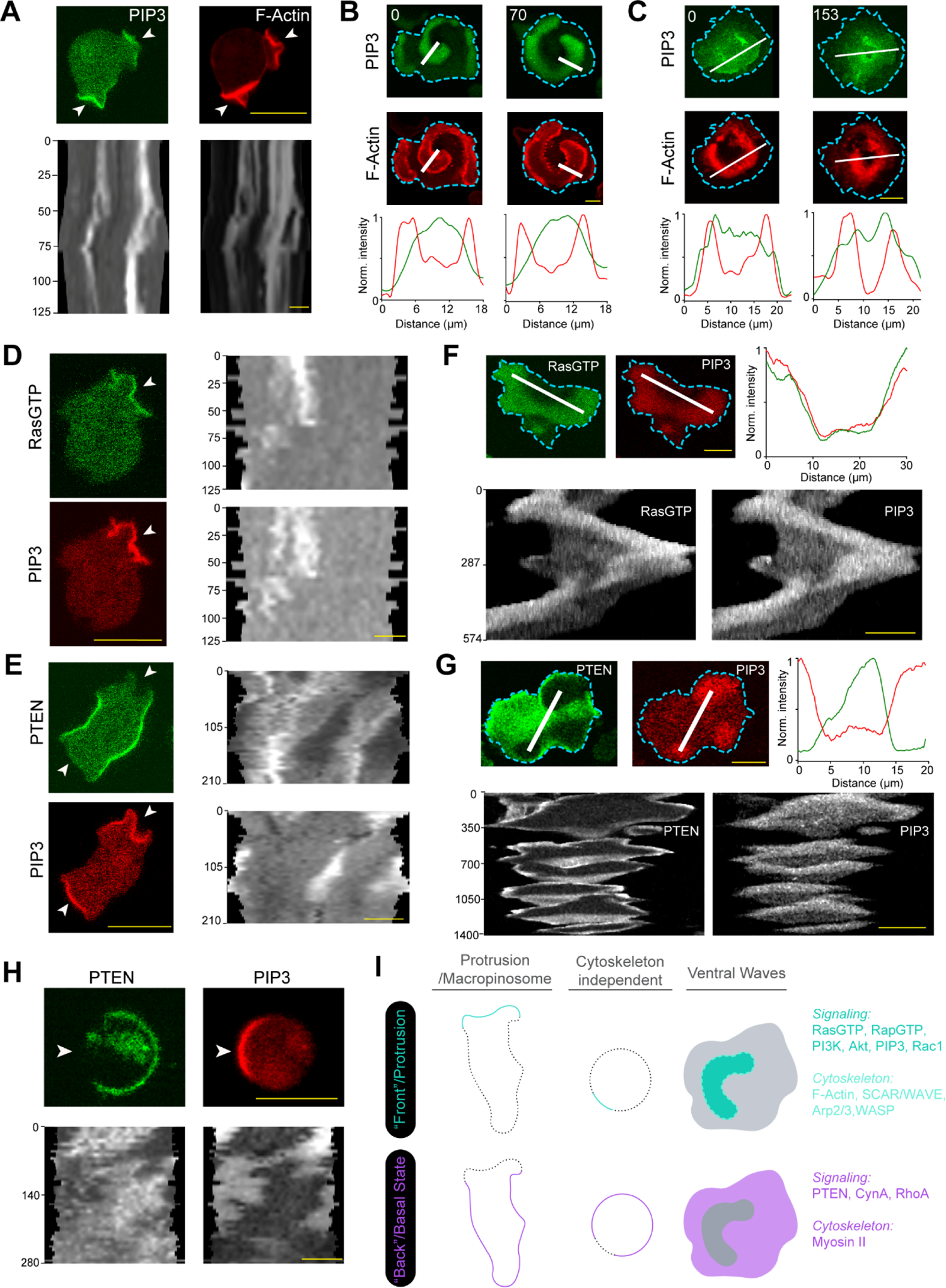
Cells generate two mutually exclusive dynamic states in the membrane during migration and ventral wave propagation. **(A)** Coordinated localization dynamics of signaling (PIP(3,4,5)P3 or PIP3) and cytoskeletal components (F-actin) in migrating *Dictyostelium* cell protrusions. PIP3 is marked by PH_crac_-GFP, newly polymerizing F-actin is marked by LimE_Δcoil_-mCherry. Top panel: Representative live-cell images (White arrows: Protrusions rich in both F-actin and PIP3). Bottom panel: 360° membrane kymographs show consistency of coordination. Here and in all other grayscale kymographs, numbers on the left denote time in seconds, unless otherwise mentioned. **(B and C)** Coordinated propagation of signaling and cytoskeletal components in ventral cortical waves of *Dictyostelium* (B) and RAW 264.7 macrophages (C). Signaling component PIP3 is marked by PH_crac_-GFP in *Dictyostelium* and by PH_AKT_-GFP in macrophages. Newly polymerizing F-actin is marked by LimE_Δcoil_-mCherry in *Dictyostelium* and by Lifeact-mCherry in macrophages. Top two panels show live-cell time-lapse images and bottom panels show line-scan intensity profile along the white lines. (**D and E**) Activated Ras (marked by Ras-Binding Domain of mammalian Raf1; RBD) and PIP3 co-localizes in the protrusions (D), whereas PTEN dissociates from the protrusions and displays a tight complementary kinetics with respect to PIP3 (E), in migrating *Dictyostelium* cells. Representative live-cell images (left) and 360° membrane kymographs (right) are shown; White arrows: Protrusions/front-states, marked by RBD and/or PIP3. **(F and G)** In propagating ventral waves of *Dictyostelium* cells, activated Ras and PIP3 dynamically colocalizes and defines the front-state regions (F), whereas PIP3 and PTEN exhibit consistent complementarity (G). Representative live-cell image, line-scan intensity profile, and line-kymographs are shown. (**H**) Complementary spatiotemporal distribution of signaling front (PIP3) and back markers (PTEN) is independent of cytoskeleton. Here *Dictyostelium* cells exhibit symmetry breaking in membrane, in presence of 5 μM Latrunculin A (inhibitor of F-actin polymerization). Representative live-cell images and 360° membrane kymograph are shown. White arrows denoting front-states. (**I**) Overall schematic shows the front-back complementary patterning in three different scenarios: migrating cell protrusions, cytoskeleton independent cortical symmetry breaking, and propagating ventral waves. When imaging first two cases, we studied a 1D profile, whereas for the case of ventral waves, we observed a 2D profile at the substrate-attached surface. A number of examples of established signaling and cytoskeletal components are listed and categorized based on their front or back-state localization. Note that, in all situations, when a front is created from the back or basal-state of the membrane, all back markers move away from that particular area, maintaining asymmetry and complementarity. For all figures, scale bars are 10 μm.

**Figure S2.**
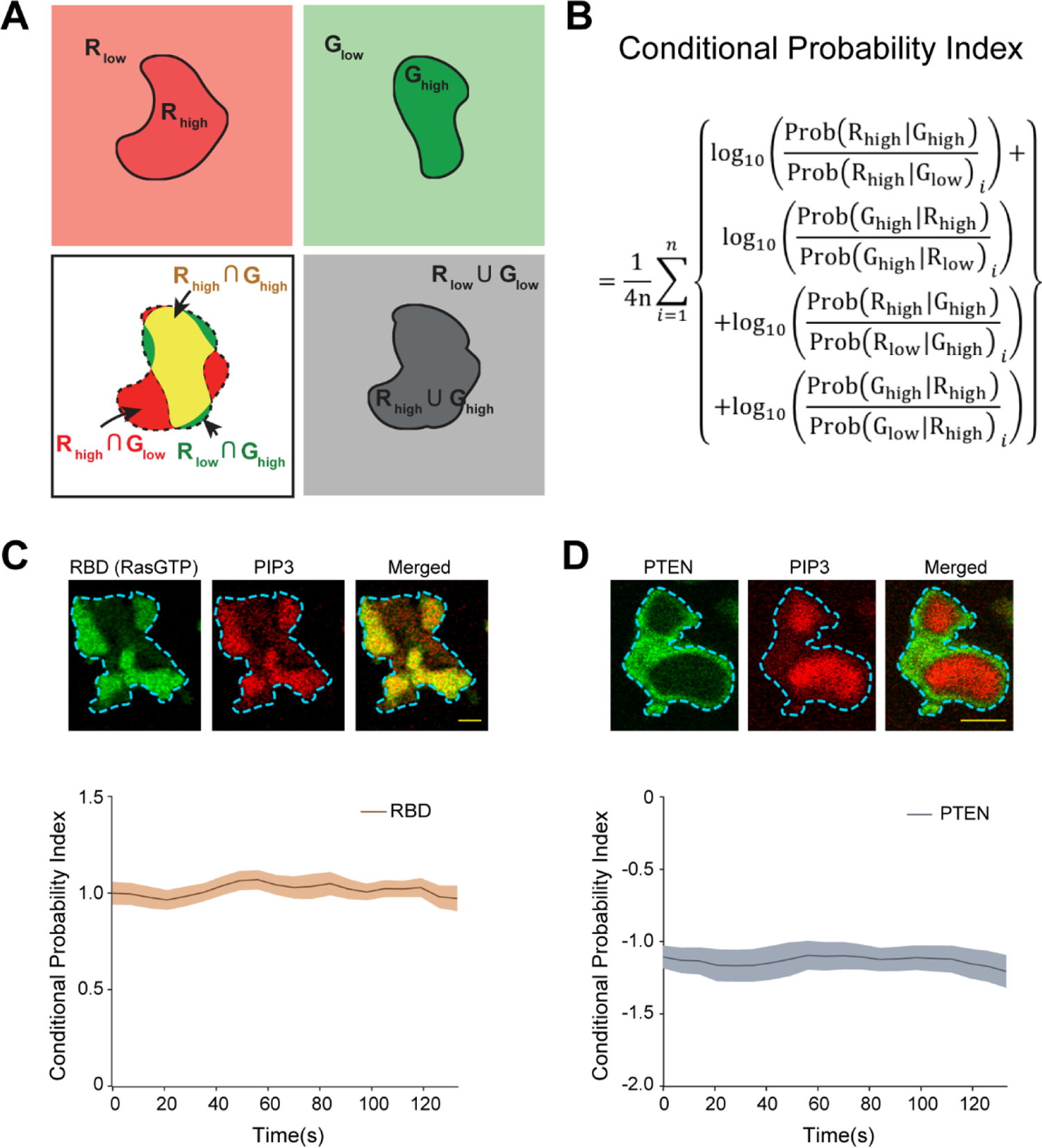
Developing conditional probability index as a metric to quantify the extent of co-localization and complementary localization. **(A)** Schematics showing the application of the concepts of conditional probability in quantifying the degree of colocalization between two entities, R and G. The regions of the high enrichments of the species R and G are denoted as R_high_ and G_high_ against the depleted states of R_low_ and G_low_, respectively (top panels). The overlapped region (yellow in the bottom left panel) denotes R_high_∩G_high_. The other necessary probabilities are also shown which are required in the computation of the respective Conditional Probability Index (CP index). **(B)** The mathematical description of the CP index. As usual, P(R_high_|G_high_) denotes Probability of selecting R_high_, given G_high_ is already selected (please see methods for details). (**C and D**) Time series plots of CP indices of established back protein PTEN (C) and established front sensor RBD (D); number of cells n_c_ =15 for RBD (C) and n_c_ =17 for PTEN (D). To generate CP index time-plots, each cell was analyzed for n_f_ =20 frames; data are mean ± SEM. Top panels show representative images of ventral waves in cells co-expressing either PH_Crac_ and RBD (C) or PH_Crac_ and PTEN (D). Note that the CP index value of PTEN is negative and RBD is positive which demonstrate their back- and front-state localization, respectively. Throughout this paper, all CP indices are calculated with respect to PIP3.

**Figure S3.**
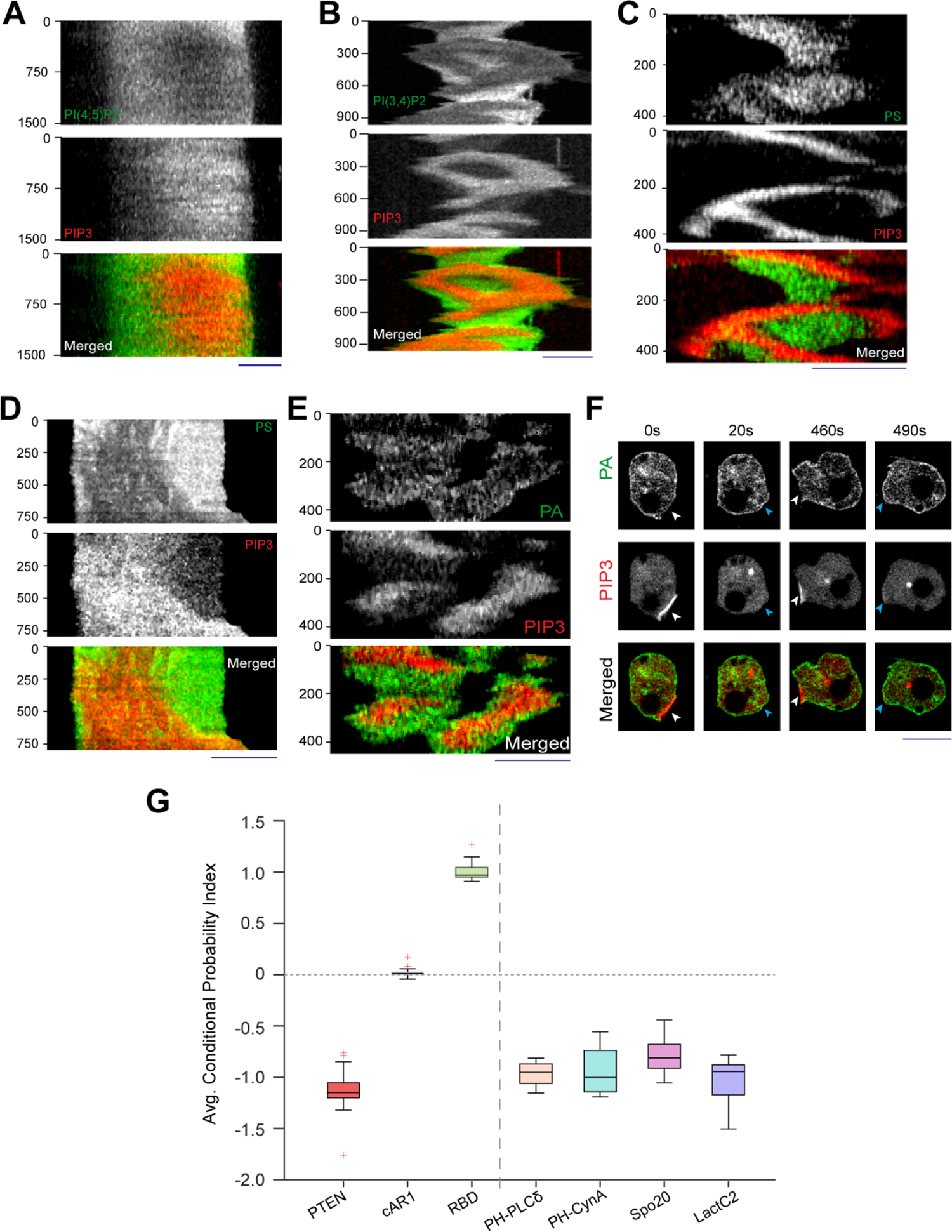
PI(4,5)P2, PI(3,4)P2, PS, and PA exhibit consistent yet dynamic back-state distribution. **(A)** Representative line-kymograph of ventral waves in RAW 264.7 macrophage cells co-expressing PI(4,5)P2 biosensor, GFP-PH_PLCδ_, and PIP3 biosensor, PH_Akt_-mCherry. Time-lapse images and line-scan intensity profiles were shown in Fig. 1c. **(B)** Representative line-kymograph of ventral waves in *Dictyostelium* cells co-expressing PI(3,4)P2 biosensor, PH_CynA_-KikGR and PIP3 biosensor, PH_Crac_-mCherry. Time-lapse images and line-scan intensity profiles were shown in Fig. 1D. **(C)** Representative line-kymographs of ventral wave pattern shown in *Dictyostelium* cells co-expressing PS sensor GFP-LactC2 and PIP3 sensor, PH_Crac_-mCherry. Time-lapse images and line-scan intensity profiles were shown in Fig. 1G. **(D)** Representative line-kymographs of ventral wave pattern in RAW 264.7 macrophage cells co-expressing PS sensor GFP-LactC2 and PIP3 sensor, PH_Akt_-mCherry. Time-lapse images and line-scan intensity profiles were shown in Fig. 1h. **(E)** Representative line-kymographs of ventral wave pattern in *Dictyostelium* cells co-expressing PA sensor, GFP-Spo20 and PIP3 sensor, PH_Crac_-mCherry. Time-lapse images and line-scan intensity profiles were shown in Fig. 1J. **(F)** Time-lapse images of migrating *Dictyostelium* cells co-expressing GFP-Spo20 and PH_crac_-mCherry. White arrows: Protrusions where PIP3 is enriched and PA is depleted. Blue arrows: Spo20 returned back to the membrane as protrusions were eventually retracted and membrane domain returned to its basal back-state. **(G)** Box and Whisker plot of time-averaged CP indices of four anionic phospholipids (PI(4,5)P2, PI(3,4)P2, PS, and PA), together with uniform membrane marker control cAR1, back protein PTEN, and front sensor RBD; n_c_=16 cells for PI(4,5)P2/PH_PLCδ_, n_c_=11 cells for PI(3,4)P2/PH_CynA_, n_c_=15 cells for PS/LactC2, n_c_=16 cells for PS/Spo20, n_c_ =20 cells for cAR1, n_c_ = 17 cells for PTEN, n_c_ = 15 cells for RBD. To generate each datapoint, n_f_ =20 frames were averaged for the above-mentioned number of cells.

**Figure S4.**
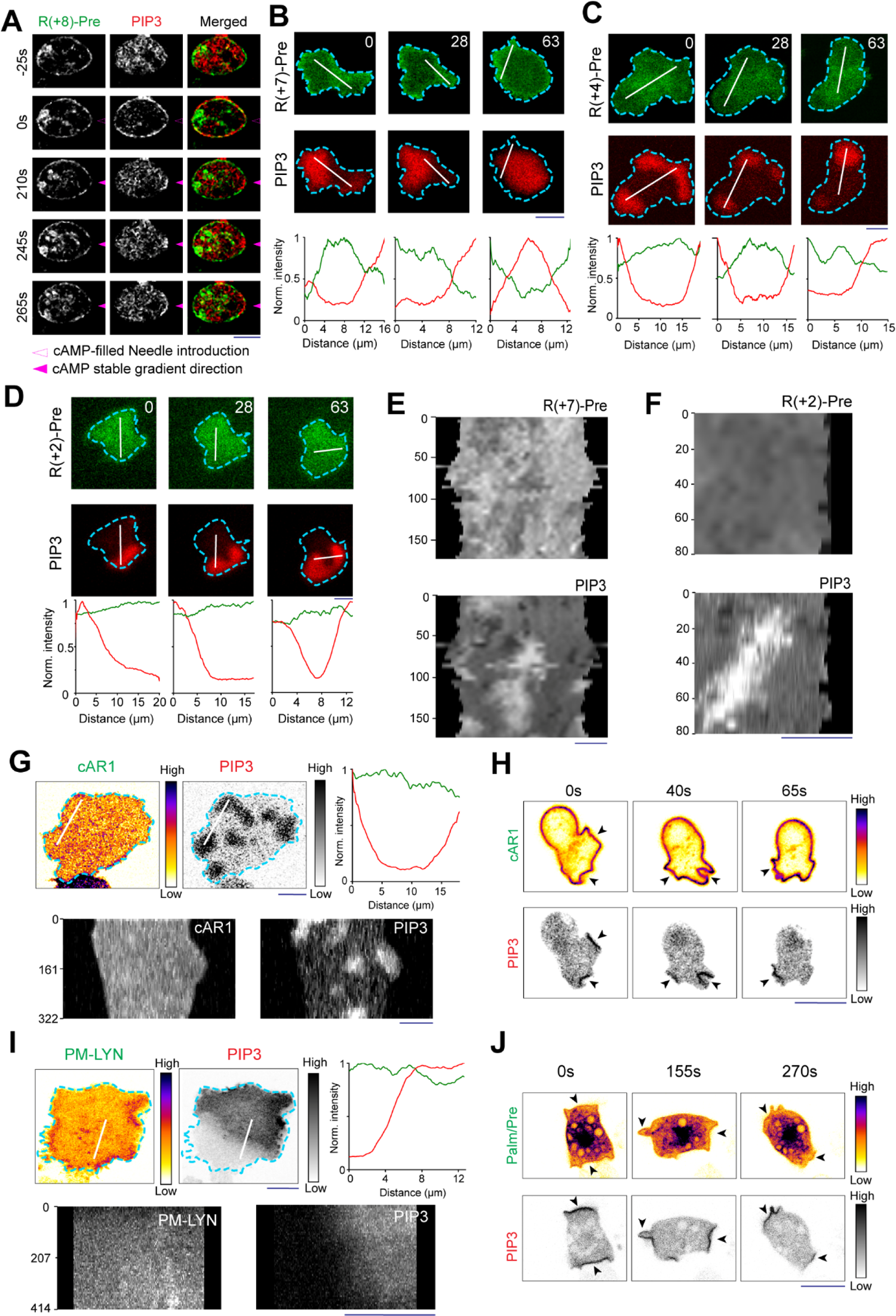
Spatiotemporal organization of different mutated charge sensors and uniform membrane controls. **(A)** Representative live cell images of Dictyostelium cells co-expressing GFP**-**R(+8)-Pre and PH_Crac_-mCherry under chemotactic gradient stimulation. Solid magenta arrowhead indicates the direction of micropipette (filled with 1 μM cAMP) for gradient stimulation. Dashed magenta arrowhead indicates the introduction of needle (t=0s) which is manifested by the transient global response in PH_Crac_ channel. Cells were pre-treated with Latrunculin A. **(B-D)** Live-cell time-lapse images and linescan intensity profiles of *Dictyostelium* cells expressing PIP3 biosensor, PH_crac_-mCherry, along with GFP-R(+7)-Pre (B) or GFP-R(+4)- Pre (C) or GFP-R(+2)-Pre (D), in ventral waves of *Dictyostelium* cells, displaying decreasing extent of back-state preference. The first time points were showed in Fig. 2J (in grayscale colormap). (**E and F**) The 360° membrane kymographs of cells shown in Fig. 2k indicates R(+7)-Pre consistently moves away from PIP3-rich protrusions (E), whereas R(+2)-Pre is uniform over the cortex (F). (**G**) Representative live-cell images, linescan intensity profiles, and representative line-kymographs of ventral waves in *Dictyostelium* cells co-expressing PH_crac_-mCherry and membrane marker cAR1-GFP, demonstrating that cAR1 does not distribute to front- or back-state regions and it is consistently uniform over the membrane. The “Fire invert” LUT was used for cAR1 so that if can clearly show any small inhomogeneity. (**H**) Representative live-cell time-lapse images of migrating *Dictyostelium* cells co-expressing PH_crac_-mCherry and cAR1-GFP shows that cAR1 is symmetric over the membrane. Black arrows: cAR1 does not move away from the PIP3-rich protrusions. (**I**) Representative live-cell images, linescan intensity profiles, and representative line-kymographs of ventral waves in RAW 264.7 cells co-expressing PH_AKT_-mCherry and membrane marker, LYN-GFP, showing consistent uniform profile of LYN over the membrane and no depletion in front-state area. (**J**) Representative live-cell time-lapse images of migrating *Dictyostelium* cells co-expressing PH_crac_-mCherry and GFP-Palm/Pre, showing a symmetric profile of Palm/Pre over the membrane. Black arrows: Protrusions/front-states.

**Figure S5.**
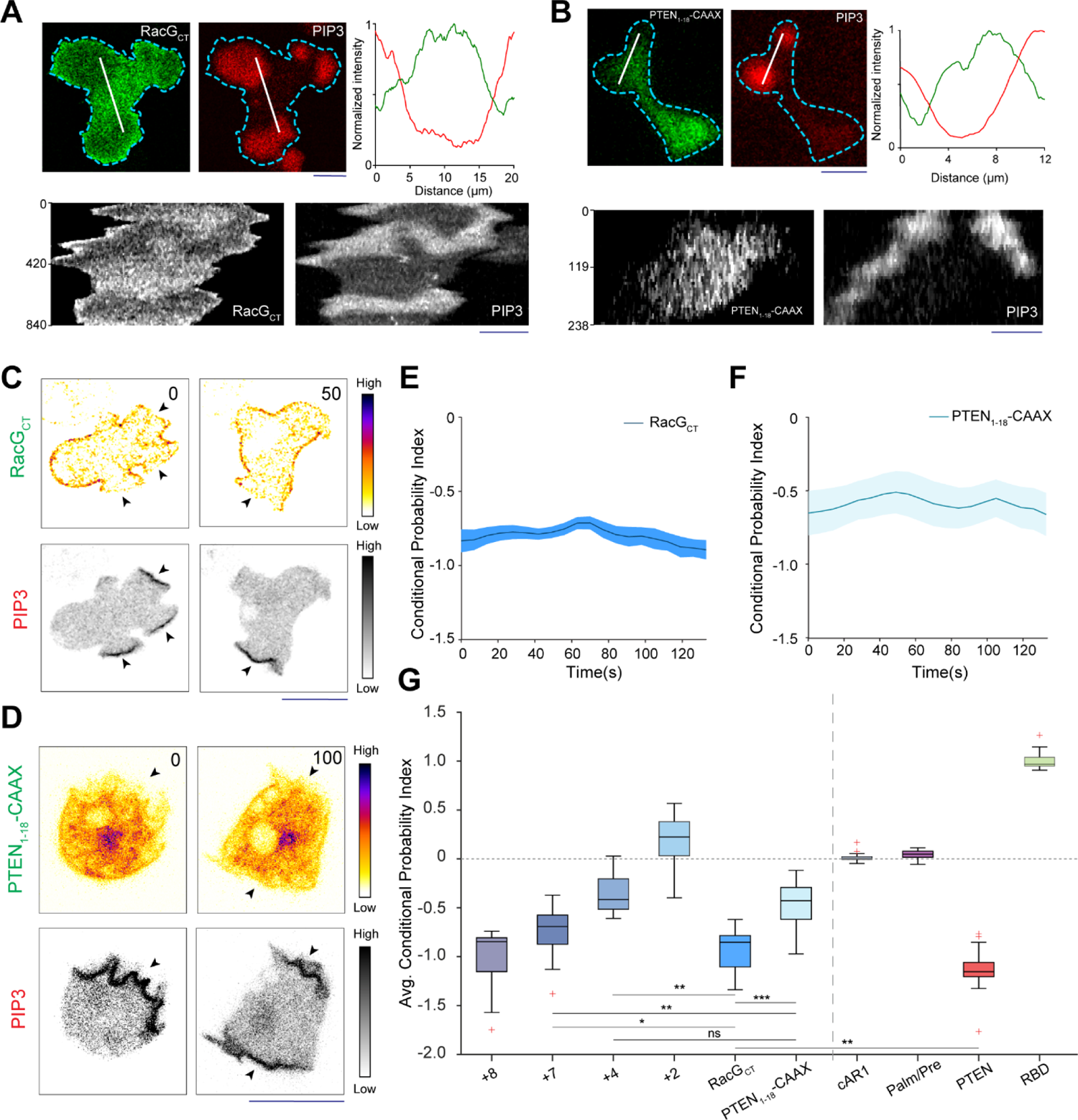
Different polybasic sequences localize to back-state regions depending on their charge, irrespective of their exact amino acid sequences. **(A and B)** Representative live-cell images, linescan intensity profiles, and representative line-kymographs of *Dictyostelium* cells co-expressing PH_Crac_-mCherry and GFP-RacG_CT_ (A) or PTEN_1-18-_CAAX (B), demonstrating consistent dynamic back distribution for RacG_CT_ and limited back distribution for PTEN_1-18_-CAAX in ventral waves. (**C and D**) Representative live-cell time-lapse images showing distribution of RacG_CT_ (C) or PTEN_1-18_-CAAX (D) in migrating *Dictyostelium* cells, demonstrating a localization profile analogous to (A and B). (**E and F**) Time series plot of CP index of RacG_CT_ (E) and PTEN_1-18_-CAAX (F) show the extent of back localization; n_c_=17 for RacG_CT_ (E), n_c_= 12 for PTEN_1-18_-CAAX (F); mean ± SEM. **(G)** Comparison of localization profile by Box and Whisker plot of time-averaged CP indices of all surface charge sensors, together with uniform membrane marker controls, back protein PTEN, and front sensor RBD; R(+8)-Pre: n_c_ =30, R(+7)-Pre: n_c_ =23, R(+4)-Pre: n_c_ =20, R(+2)-Pre: n_c_ =12, RacG_CT_: n_c_ =17, PTEN_1-18_-CAAX: n_c_ =12, cAR1: n_c_ =20, Palm/Pre: n_c_ =11, PTEN: n_c_ =17, RBD: n_c_ =15. To generate each datapoint, n_f_ =20 frames were averaged for mentioned number of cells (n_c_). Box and whiskers are graphed as per Tukey’s method. All p-values by Mann-Whitney-Wilcoxon test.

**Figure S6.**
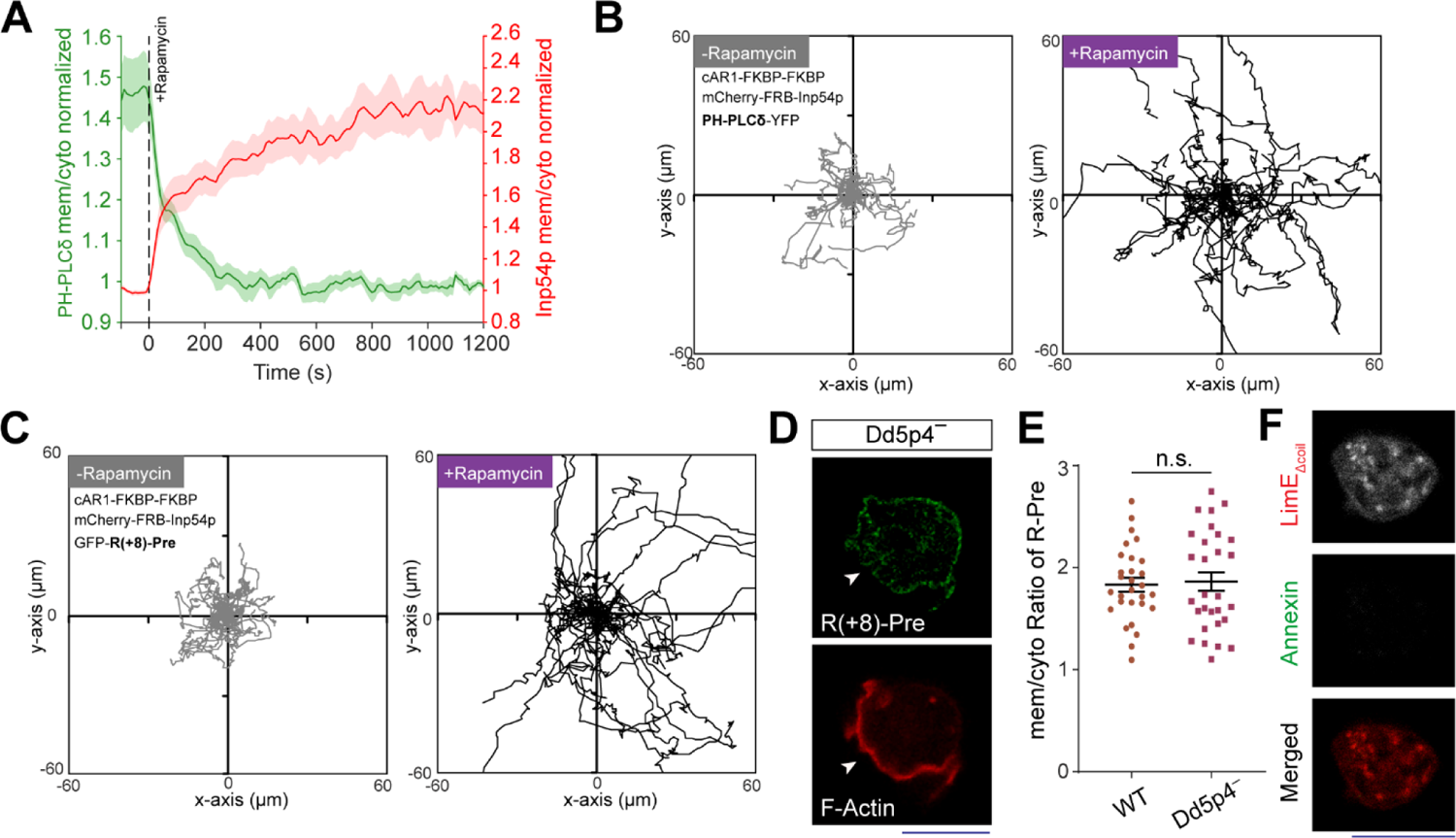
Dynamics of surface charge sensor in PI(4,5)P2 and PI(3,4)P2 depleted cells. **(A)** Time course of membrane/cytosol ratio of PH_PLCδ_ and Inp54p upon rapamycin addition (indicated by black dashed vertical line), in *Dictyostelium* cells co-expressing cAR1-FKBP-FKBP, mCherry-FBP-Inp54p, and PH_PLCδ_-GFP, demonstrating PH_PLCδ_ dissociated from membrane upon PI(4,5)P2 depletion; n=17 cells; mean ± SEM. **(B and C)** Cell tracks show the migration profile of *Dictyostelium* cells expressing Chemically induced dimerization system cAR1-FKBP-FKBP and mCherry-FBP-Inp54p, along with GFP-R(+8)-Pre (B) or PH_PLCδ_-GFP (C), before and after rapamycin induced recruitment. Tracks demonstrating similar change in migration profile in both cases, as quantified in terms of migration speed in Figure 3D. To generate each track for n_c_=32 cells (in each case), cells were followed for n_f_=60 frames (7s/frame). **(D)** Representative image of Dd5p4^-^ *Dictyostelium* cell (where PI(3,4)P2 level is low) co-expressing GFP-R(+8)-Pre and LimE_Δcoil_-mRFP displaying characteristic membrane association and back localization of R(+8)-Pre; white arrows denote F-actin rich protrusions. **(E)** Quantification of membrane association of R(+8)-Pre in wild type and Dd5p4-cells, in terms of membrane/cytosol ratio; n=29 cells in each case; p-value by Mann-Whitney-Wilcoxon test. **(F)** Example of a quiet or non-protrusion forming *Dictyostelium* cells expressing LimE_Δcoil_-GFP, whose outer leaflet of membrane was allowed to transiently bind with Annexin V.

**Figure S7.**
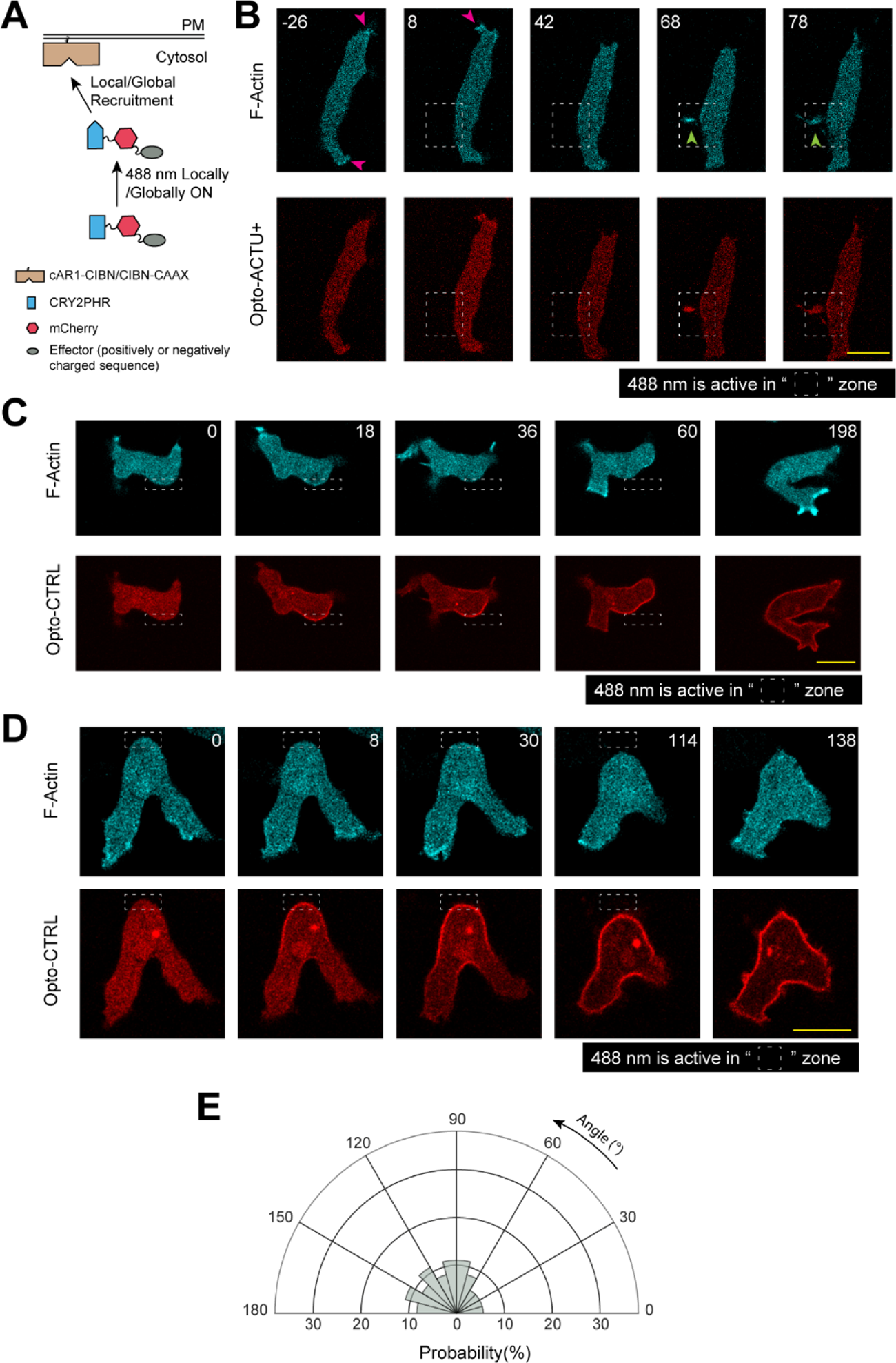
Local recruitment of Opto-ACTU+ in polarized cells generates protrusion *de novo* near recruitment area, whereas uncharged control Opto-CTRL recruitment cannot elicit any change. **(A)** Scheme for optogenetic recruitment. Turning on 488 nm laser changes the conformation of CRY2PHR module and as a result it gets recruited to the plasma membrane-bound CIBN. The cAR1-CIBN is used in all *Dictyostelium* systems and CIBN-CAAX is used in all mammalian systems as membrane anchor. **(B)** A representative example of *de novo* formation of protrusion from a position of choice in the back-state region of the membrane by spatially confined recruitment of Opto-ACTU+. Magenta arrows: old protrusions, Green arrows: new protrusions. The experimental setup and another example are shown in Fig. 4c and 4d. **(C and D)** Two representative examples of spatially confined optogenetic recruitment of Opto-CTRL demonstrating no increase in protrusion generation from the site of recruitment. In (B-D), the numbers on the images denote time in seconds. **(E)** Polar histogram show probability of protrusion formation at the different parts of the cortex with respect to Opto-CTRL recruitment area; n_c_=20 cells, n_p_=34 protrusions.

**Figure S8.**
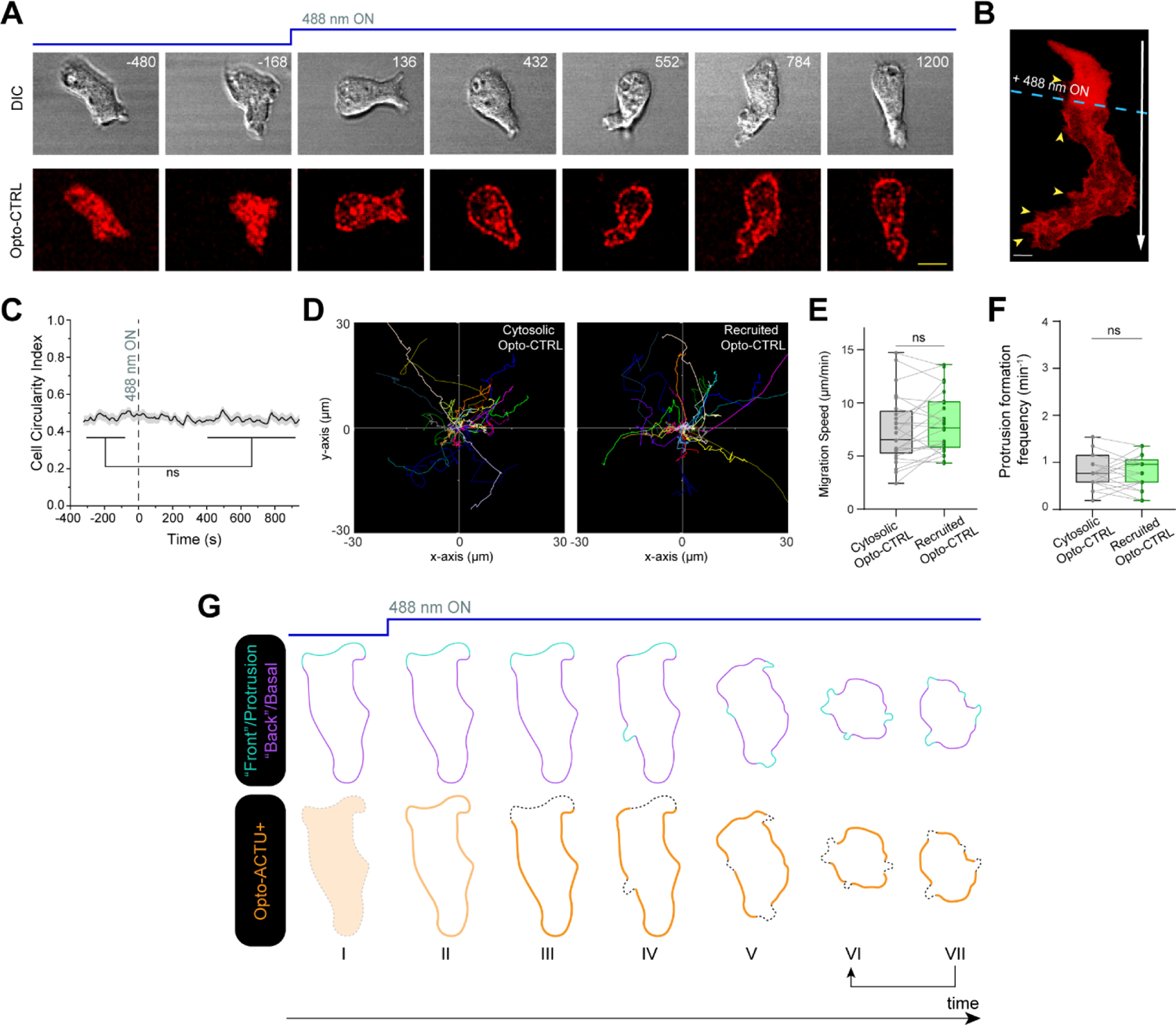
Unlike Opto-ACTU+, recruited uncharged Opto-CTRL does not exhibit any preferential localization and cannot induce any phenotypic changes in terms of migration, polarity, or protrusion formation. **(A and B)** Time-lapse snapshots (A) and time-stack (B) demonstrate the unaltered cell morphology and migration behavior in polarized *Dictyostelium* cells upon recruitment of Opto-CTRL. Numbers are time in seconds (A). 488 nm switched ON globally at t=0s. Yellow arrows: Opto-CTRL is uniform over cortex and did not move away from protrusions (B). **(C-F)** Quantification of cell morphology and migration mode in terms of cell circularity index (C), cell tracks (D), migration speed (E), and new protrusion formation frequency (F), upon Opto-CTRL recruitment (n=25 cells). Data shown as mean ± SEM over time in (C). In (D-F), for either before or after recruitment tracks, each cell tracked for n_f_=40 frames (t=320 s). Tracks were reset to the same origin in (D). For pairwise comparison, tracks are color-coded in (d), data from same cell are connected by gray line in (E) and (F). **(G)** Schematic proposing how Opto-ACTU+ recruitment changes cell morphology and migratory mode. Opto-ACTU+ is recruited globally as expected (I to II); however, presumably due to its positive charge, it quickly accumulates along the back regions of the cell (III). Consequently, new protrusions are elicited from these back regions and the cell begins to lose polarity (IV and V). And at the same time, as some areas of erstwhile back regions are converted to front, Opto-ACTU+ redistributes again to the newly formed back regions (IV to VII). This in turn generates fresh protrusions there and this entire cycle is repeated (shown in arrows between VI and VII). As a result of this series of events, protrusions are generated randomly, migration becomes impaired, and pre-existing polarity is abrogated.

**Figure S9.**
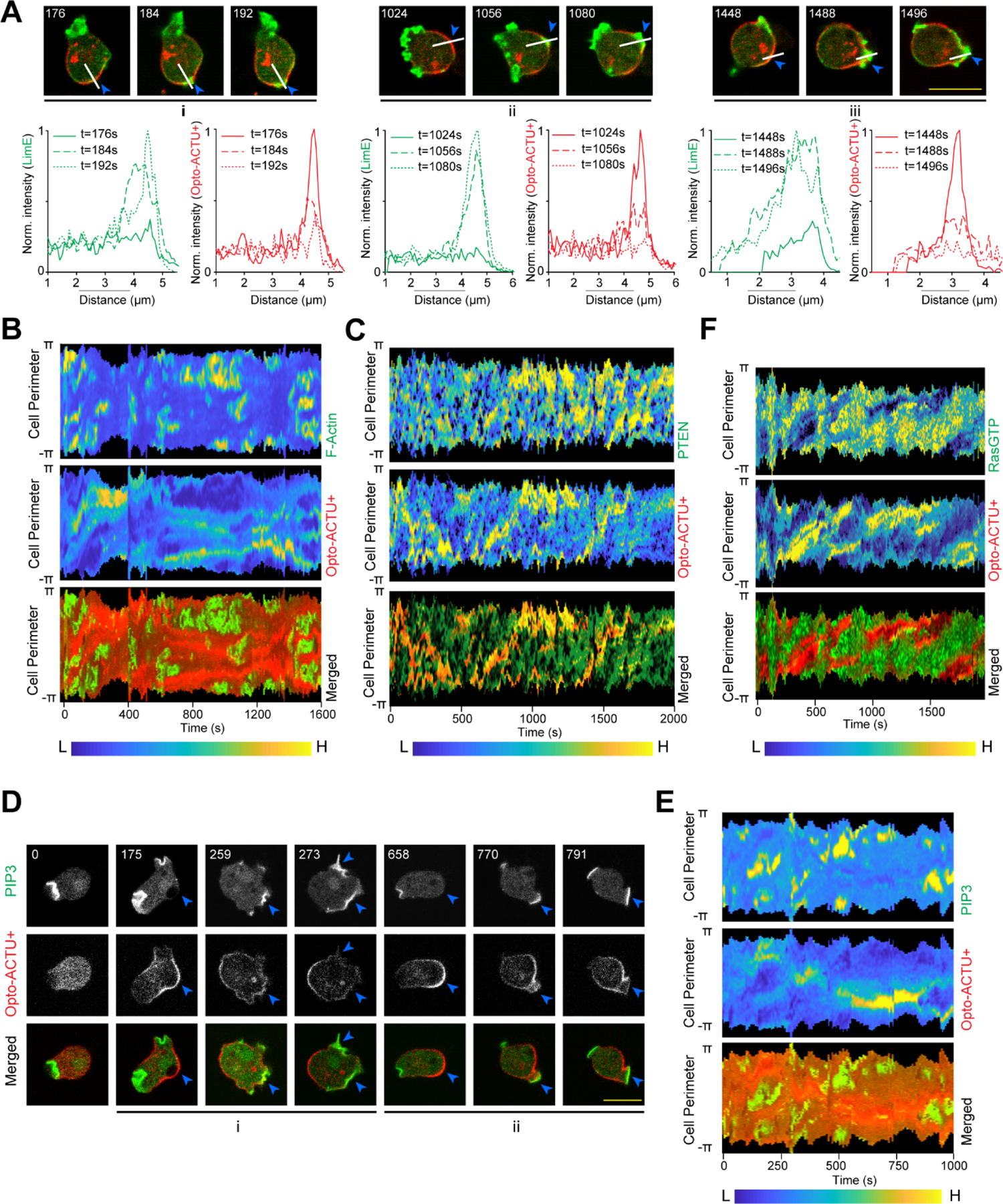
Global recruitment of Opto-ACTU+ can cause spatiotemporally confined activation of Ras/PI3K/Akt/TORC2/F-actin network components. **(A)** Intensity profiles of newly polymerized F-actin biosensor LimE-GFP and Opto-ACTU+ along the white lines (the images are same as shown in Fig. 5b) demonstrate that F-actin polymerizes in the domains of membrane where Opto-ACTU+ accumulates and when that leads to a protrusion, Opto-ACTU+ moves away with a short time delay. **(B)** 360° membrane kymograph of cell shown in Fig. 5B. (C) 360° membrane kymograph of cell shown in Fig. 5C. **(D)** 360° membrane kymograph of cell shown in Fig. 5D. **(E)** Time-lapse live cell images of *Dictyostelium* cells co-expressing Opto-ACTU+, cAR1-CIBN, and PIP3 biosensor PH_Crac_ where recruitment was started at t=0s. Numbers show time in seconds. The “i” and “ii” are showing two different PIP3 production events which eventually lead protrusion formation. For each event, blue arrowheads in are showing the areas where Opto-ACTU+ was first accumulated which in turn became the areas of PIP3 production and eventually after protrusion formation, Opto-ACTU+ moved away to a newer back-state area. **(F)** 360° membrane kymograph of cell shown in (E).

**Figure S10.**
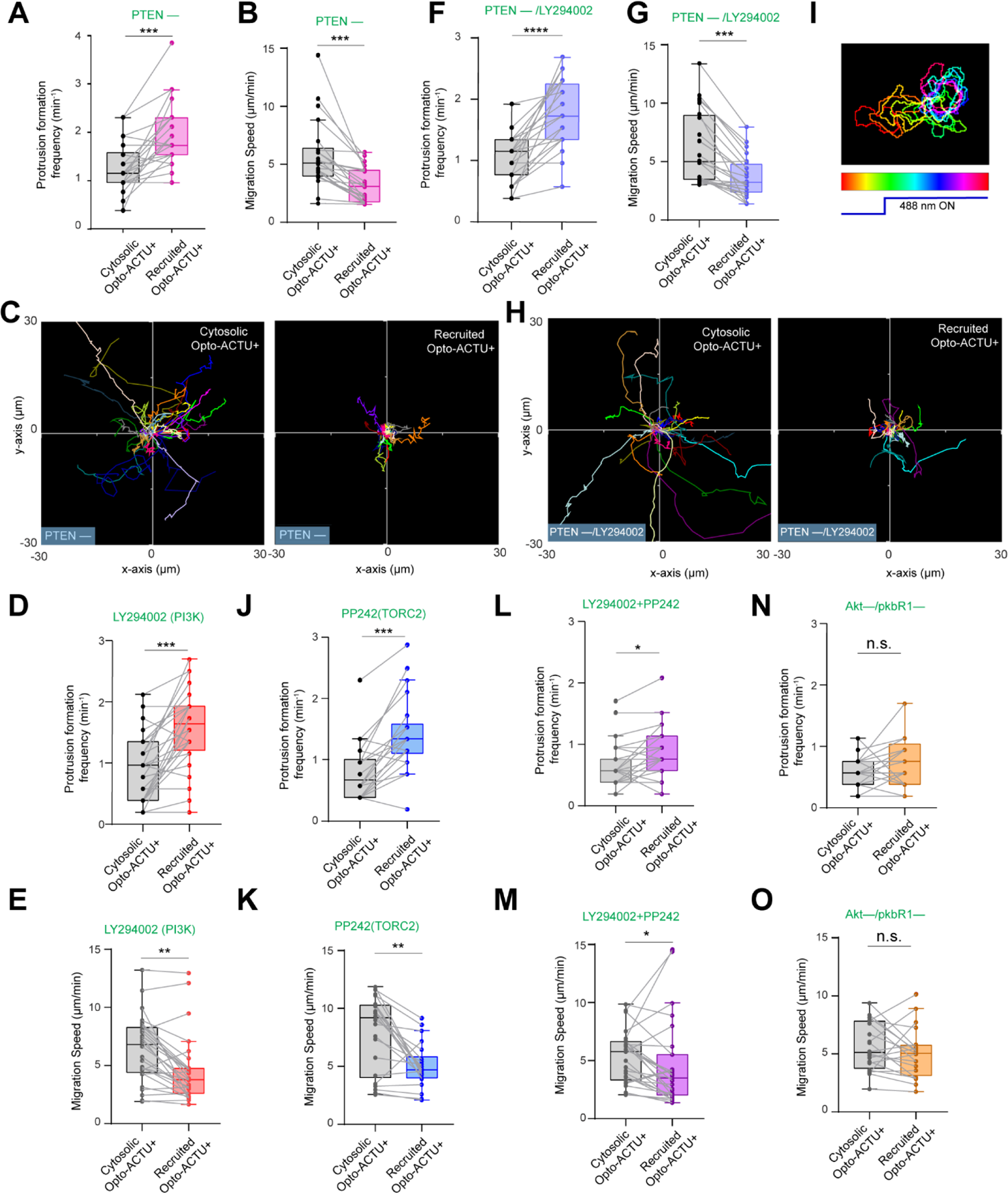
The effect of specific genetic and pharmacological inhibitions upon the phenotypic changes induced by Opto-ACTU+ recruitment. **(A-C)** Quantification of migration profile changes in terms of new protrusion formation frequency (A), speed (B), and cell tracks (C), upon Opto-ACTU+ recruitment, in PTEN– *Dictyostelium* cells; n_c_= 22 cells. **(D, E)** Quantification of migration profile changes in terms of new protrusion formation frequency (D) and speed (E), upon Opto-ACTU+ recruitment in *Dictyostelium* cells, pre-treated with PI3K inhibitor LY294002; n_c_= 28 cells. **(F-H)** Quantification of migration profile changes in terms of new protrusion formation frequency (F), speed (F), and cell tracks (H), upon Opto-ACTU+ recruitment in PTEN– *Dictyostelium* cells, pre-treated with PI3K inhibitor LY294002; n_c_= 24 cells. **(I)** Temporally color-coded cell outlines of a representative migrating PTEN– *Dictyostelium* cells, pre-treated with PI3K inhibitor LY294002, showing cell morphology and migratory profile before and after 488nm was turned on to recruit Opto-ACTU+ (corresponding to Video S11). **(J-O)** Quantification of migration profile changes in terms of new protrusion formation frequency (J, L, N) and migration speed (K, M, O) upon Opto-ACTU+ recruitment under different genetic and pharmacological inhibitions. In (J-K) cells were pre-treated with PP242 to inhibit TORC2 (n_c_= 22 cells); in (L-M) cells were pre-treated with both LY294002 and PP242 to simultaneously block PI3K and PP242 (n_c_= 27 cells); in (K-L), Akt^—^/PKBR1^—^ double knockout cell line was used (n_c_= 21 cells). For each case, each of the n_c_ cells were tracked for n_f_=40 frames (8 sec/frame was imaging frequency) and time averages were taken. Tracks were reset to the same origin in (C) and (H). For pairwise comparison, tracks are color-coded in (C) and (H). In all box plots here, for pairwise comparison, data from same cell are connected by gray lines. The p-values by Mann-Whitney-Wilcoxon test.

**Figure S11.**
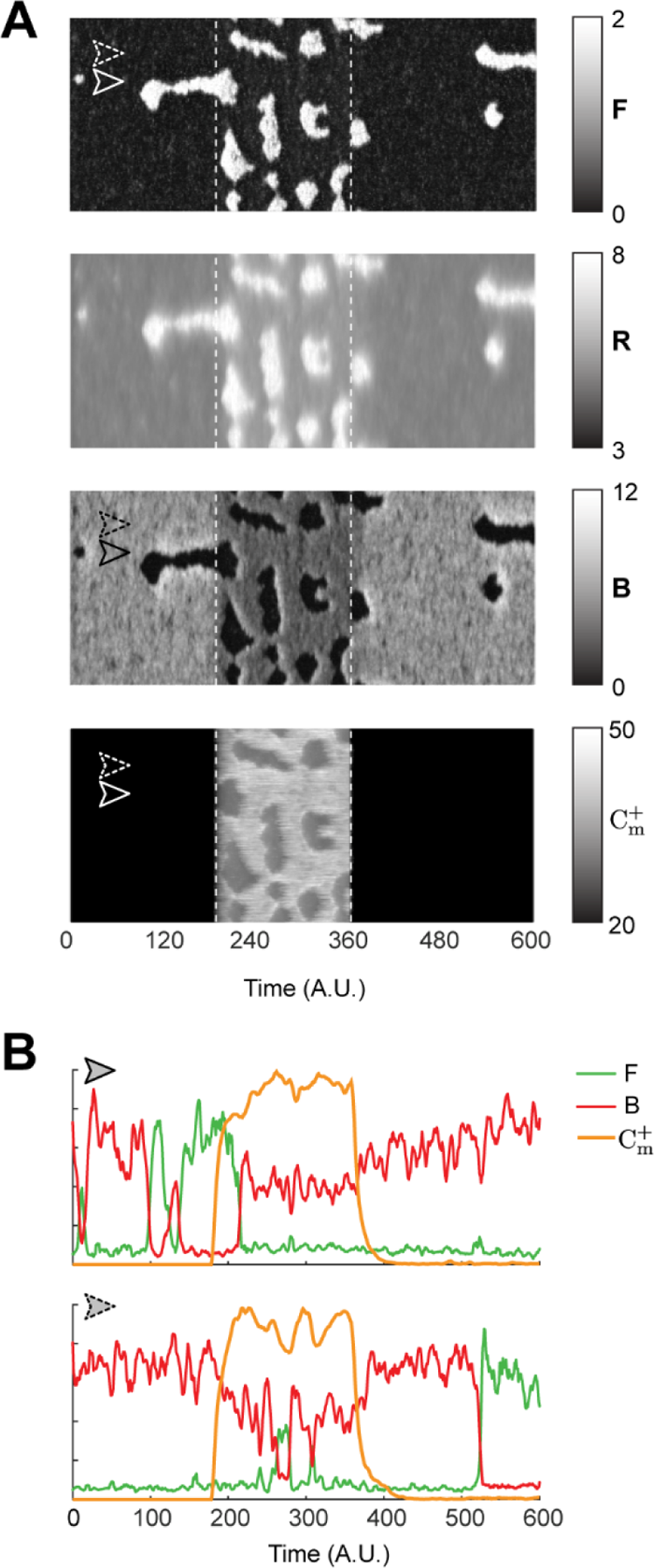
In silico reversible recruitment of Opto-ACTU+ can recreate the reversibility of polarity breaking and protrusion formation events that was observed experimentally. **(A)** The simulated kymographs of F (first), R (second), B (third) and *C_m_*^+^ (fourth) in response to global reversible recruitment. The instant of recruitment is shown by the first white dashed line and recruitment was stopped in the second dashed line. **(B)** Line scans at two locations (denoted by dashed and solid arrows) on the simulated kymographs showing the temporal profiles of F (green), B (red) and *C_m_*^+^ (orange).

**Figure S12.**
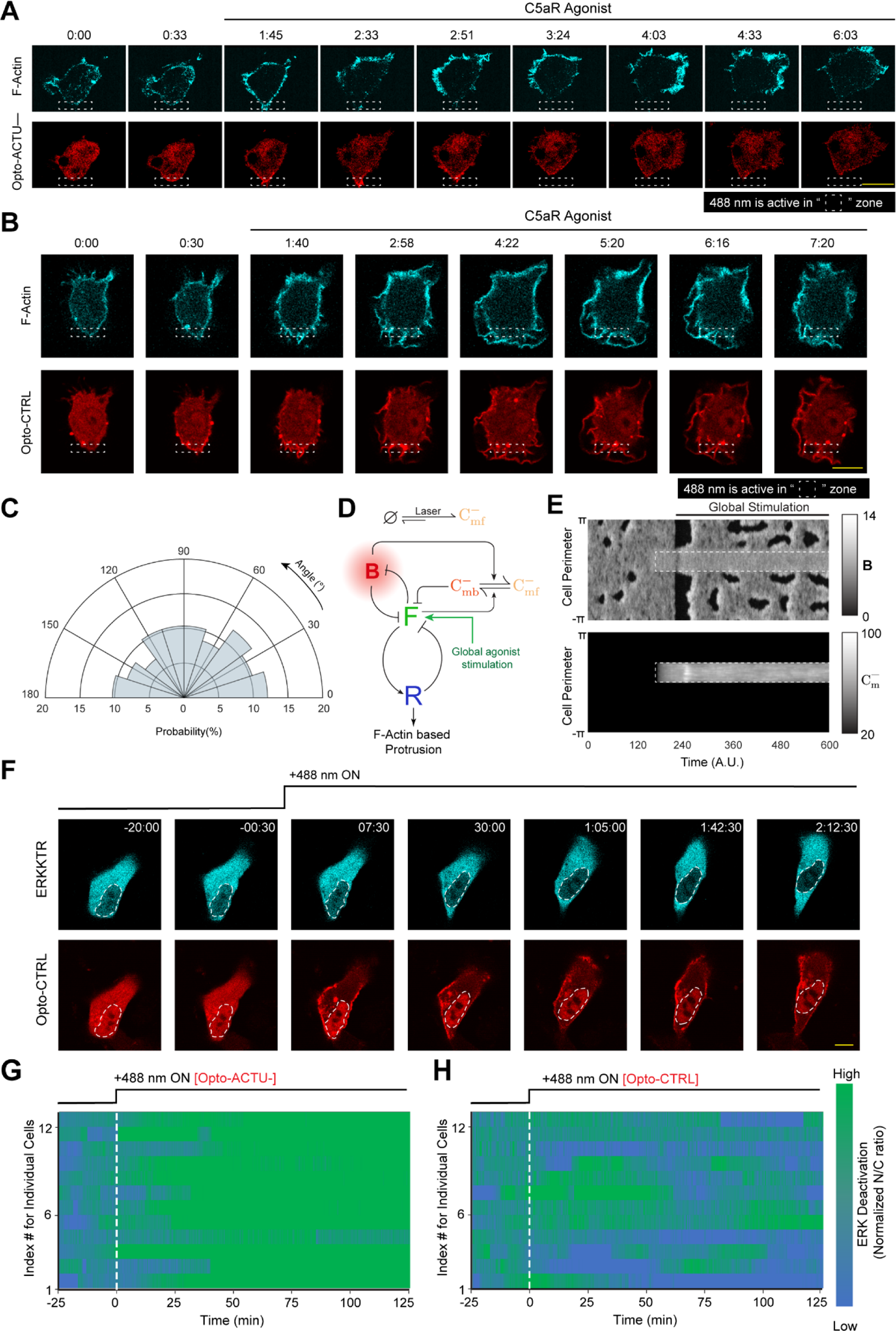
Selective recruitment of uncharged control Opto-CTRL cannot suppress protrusion in RAW 264.7 macrophages or deactivate ERK in MCF10A cells. **(A)** Representative live-cell time-lapse images of RAW 264.7 cells undergoing light-triggered spatially confined recruitment of Opto-CTRL, followed by global stimulation by C5a receptor agonist, demonstrating selective protrusion suppression in the site where Opto-ACTU- was locally recruited and robust protrusion formation in other areas of cortex. Time in min:sec format. Cells were co-expressing Opto-ACTU–, CIBN-CAAX, and Lifeact-mVenus. **(B)** Representative live-cell time-lapse images of RAW 264.7 cells undergoing light-triggered spatially confined recruitment of Opto-CTRL, followed by global stimulation by C5a receptor agonist. Time in min:sec format. Cells were co-expressing Opto-CTRL, CIBN-CAAX, and Lifeact-mVenus. **(C)** Polar histogram indicating probability of protrusion formation is essentially uniform over the cortex; n_c_=12 cells, n_p_=59 protrusions. **(D)** Schematic showing coupled system of excitable network, polarity module and Opto-ACTU-system along with global agonist stimulation. **(E)** The simulated kymographs of B (top) and *C_m_*^−^ (bottom) in response to local recruitment of Opto-ACTU-. The location of recruitment is denoted by the white dashed box. The solid black line denotes the span of global agonist stimulation. **(F)** Representative live-cell time-lapse images of a MCF10A cell displaying ERKKTR maintaining its cytosolic distribution upon Opto-CTRL recruitment, demonstrating no substantial ERK deactivation; cells were pre-treated with and maintained in a saturated dose of EGF throughout the experiment. Time in hr:min:sec format; 488 nm laser was first turned ON at t=0 min. **(G and H)** Individual cell level changes in the nuclear/cytosolic ratio of ERKKTR over time, upon recruitment of Opto-ACTU-(G) or Opto-CTRL (H). Population average is in Figure 7I. The color scale shown in right is applicable to both panels.

**Table S1.**
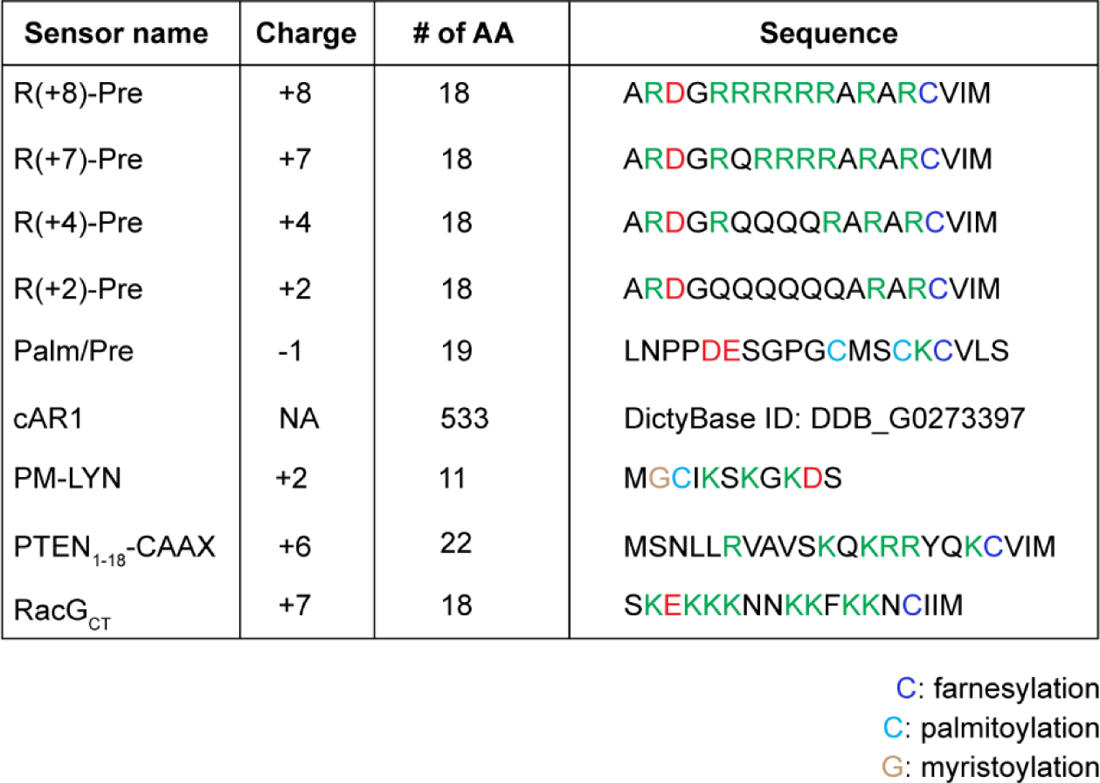
Different charge sensors and membrane marker controls. For each one, the net charge, sequence length, and exact sequence or reference is listed. Green color denotes positively charged amino acids and red color denotes negatively charged amino acids.

**Table S2.**
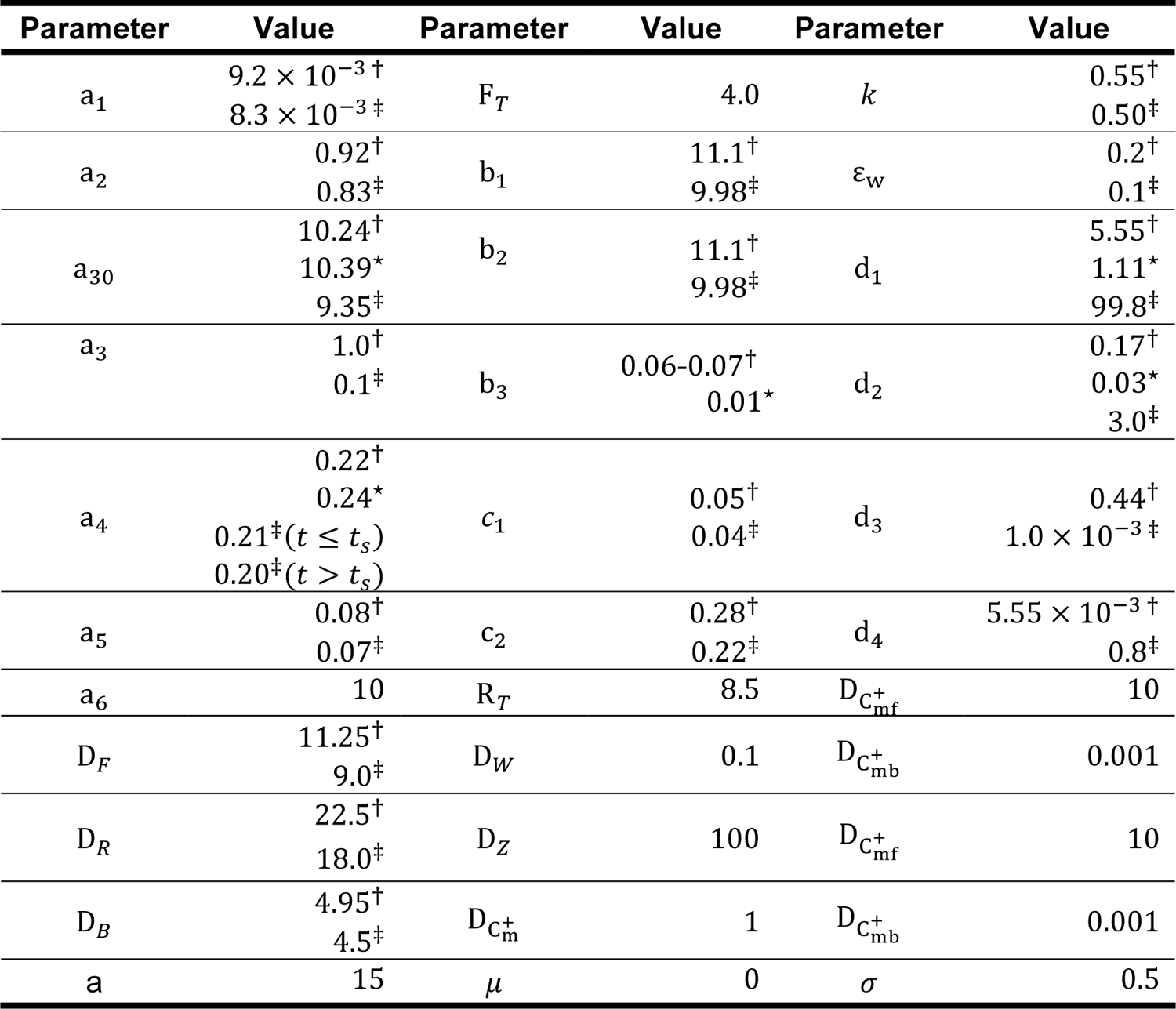
Parameter values for the computational model.

| Parameter | Value | Parameter | Value | Parameter | Value |
| --- | --- | --- | --- | --- | --- |
| $a_1$ | $9.2 \times 10^{-3}^\dagger$<br>$8.3 \times 10^{-3}^\ddagger$ | $F_T$ | 4.0 | $k$ | $0.55^\dagger$<br>$0.50^\ddagger$ |
| $a_2$ | $0.92^\dagger$<br>$0.83^\ddagger$ | $b_1$ | $11.1^\dagger$<br>$9.98^\ddagger$ | $\varepsilon_w$ | $0.2^\dagger$<br>$0.1^\ddagger$ |
| $a_{30}$ | $10.24^\dagger$<br>$10.39^*$<br>$9.35^\ddagger$ | $b_2$ | $11.1^\dagger$<br>$9.98^\ddagger$ | $d_1$ | $5.55^\dagger$<br>$1.11^*$<br>$99.8^\ddagger$ |
| $a_3$ | $1.0^\dagger$<br>$0.1^\ddagger$ | $b_3$ | $0.06-0.07^\dagger$<br>$0.01^*$ | $d_2$ | $0.17^\dagger$<br>$0.03^*$<br>$3.0^\ddagger$ |
| $a_4$ | $0.22^\dagger$<br>$0.24^*$<br>$0.21^\ddagger(t \leq t_s)$<br>$0.20^\ddagger(t > t_s)$ | $c_1$ | $0.05^\dagger$<br>$0.04^\ddagger$ | $d_3$ | $0.44^\dagger$<br>$1.0 \times 10^{-3}^\ddagger$ |
| $a_5$ | $0.08^\dagger$<br>$0.07^\ddagger$ | $c_2$ | $0.28^\dagger$<br>$0.22^\ddagger$ | $d_4$ | $5.55 \times 10^{-3}^\dagger$<br>$0.8^\ddagger$ |
| $a_6$ | 10 | $R_T$ | 8.5 | $D_{C_{mf}^+}$ | 10 |
| $D_F$ | $11.25^\dagger$<br>$9.0^\ddagger$ | $D_W$ | 0.1 | $D_{C_{mb}^+}$ | 0.001 |
| $D_R$ | $22.5^\dagger$<br>$18.0^\ddagger$ | $D_Z$ | 100 | $D_{C_{mf}^+}$ | 10 |
| $D_B$ | $4.95^\dagger$<br>$4.5^\ddagger$ | $D_{C_m^+}$ | 1 | $D_{C_{mb}^+}$ | 0.001 |
| $a$ | 15 | $\mu$ | 0 | $\sigma$ | 0.5 |

## Supplementary Video Legends

**Video S1. Dynamic localization of PI(4,5)P2 to the back-state of the membrane in both *Dictyostelium* and RAW 264.7 macrophages.** The spatiotemporal complementary patterns were observed during protrusion formation in *Dictyostelium* (A), ventral wave propagation in *Dictyostelium* (B), and ventral wave propagation in RAW 264.7 macrophages (C). Left panels show PH_PLCδ_-GFP, middle panels show PH_Crac_-mCherry (in A and B) or PH_Akt_-mCherry (in C), and the right panels show the merged view. Top left corners show time in mm:ss format.

**Video S2. During ventral wave propagation in *Dictyostelium*, PI(3,4)P2 dynamically localizes to the back-state of the membrane.** Left panels show PH_CynA_-KikGR, middle panels show PH_crac_-mCherry, and the right panels show the merged view. Top left corners show time in mm:ss format.

**Video S3. Dynamic preferential back-state distribution of phosphatidylserine in membrane during ventral wave propagation in *Dictyostelium* and RAW 264.7 macrophages.** (A) Ventral wave patterns in *Dictyostelium*. Left panel shows GFP-LactC2, middle panel shows PH_Crac_-mCherry, and the right panel shows the merged view. (B) Ventral wave propagation in RAW 264.7 macrophages. Left panels show overall pattern in the cell; right panels show the pattern in the zoomed-in area corresponding to the white rectangular boxes in the left channel. In either left or right panels, top channels show GFP-LactC2 and bottom channels show PH_Akt_-mCherry. Scale bar is 10 μm. In both (A) and (B), Top left corners show time in mm:ss format.

**Video S4. Dynamic preferential distribution of generic negative surface charge sensor to the back-state of the membrane in *Dictyostelium*.** The spatiotemporal patterns were observed during ventral wave propagation (A) and protrusion formation (B). Left panels show GFP-R(+8)-Pre, middle panels show PH_crac_-mCherry, and the right panels show the merged view. Top left corners show time in mm:ss format.

**Video S5. Dynamic preferential distribution of generic negative surface charge sensor to the back-state of the membrane in ventral waves of RAW 264.7 macrophages.** Left panel shows GFP-R(+8)-Pre, middle panel shows PH_Akt_-mCherry, and the right panel shows the merged view. Top left corners showing time in mm:ss format.

**Video S6. Dissociation of PI4P sensor PHOSH2X2 and PI(4,5)P2 sensor PHPLCδ upon recruitment of pseudojanin by chemically induced dimerization system in RAW 264.7 macrophages.** (A and B) Left panels: GFP-PHOSH2X2, Right Panels: Pseudojanin. (C and D) Left panels: GFP-PHPLCδ, Right Panels: Pseudojanin. All RAW 264.7 cells are expressing Lyn-FRB as well. In all cases, rapamycin addition time was indicated by the appearance of white-colored text “+Rapamycin” in the top middle. Top left corners showing time in mm:ss format.

**Video S7. Profile of membrane surface charge sensor R(+8)-Pre, upon recruitment of pseudojanin by chemically induced dimerization system in RAW 264.7 macrophages.** (A and B). Two examples of Pseudojanin recruitment in RAW 264.7 cells co-expressing GFP-R(+8)-Pre and Lyn-FRB. Left Panels: GFP-R(+8)-Pre; Right panels: Pseudojanin. Rapamycin addition time was indicated by the appearance of white-colored text “+Rapamycin” in the top middle. Top left corners showing time in mm:ss format.

**Video S8. Optically confined recruitment of Opto-ACTU+ can trigger protrusions *de novo*.** In (A-D), 488 nm laser was selectively irradiated inside the white rectangular boxes; top left corners showing time in seconds; untagged cAR1-CIBN was expressed. (A and B) Two examples of Opto-ACTU+ recruitment into the back-state regions of the membrane showing the generation of new protrusions in the vicinity of the recruitment. In (A) stage was moved to start recruitment at time 0s. Left panels: LimE_Δcoil_ (newly-polymerized F-Actin sensor), right panels: Opto-ACTU+. (C and D) Two examples of Opto-CTRL recruitment in the back-states of the membrane, showing no generation of new protrusions from the recruitment areas. Left panels: LimE_Δcoil_ (newly-polymerized F-Actin sensor), right panels: Opto-CTRL.

**Video S9. Global recruitment of Opto-ACTU+ to the membrane in polarized *Dictyostelium* cell elicits random protrusions and abrogates pre-existing polarity and persistent migration.** (A) Global optical recruitment of Opto-ACTU+. Left panel: Opto-ACTU+, right panel: DIC. (B) Global optical recruitment of Opto-CTRL. Left panel: Opto-CTRL, right panel: DIC. For both (A) and (B): Untagged cAR1-CIBN was expressed. Top left corners show time in seconds. The time before recruitment shown with a negative sign. The 488 nm laser was turned ON at 0s to initiate the recruitment, as depicted by the appearance of green-colored text “488 nm ON” in the bottom middle.

**Video S10. Cell morphology and migration mode changes upon global recruitment of Opto-ACTU+ are reversible.** (A) and (B) demonstrate two examples where Opto-ACTU+ was first recruited to the membrane by turning on 488 nm laser (recruitment initiated when white-colored text “488 nm OFF” switched to green-colored text “488 nm ON” in the videos) which resulted increased protrusion formation, loss of polarity, and impaired migration; then to reverse the process, the laser was turned off (when the green-colored text “488 nm ON” switched to white colored text “488 nm OFF”), Opto-ACTU+ returned to cytosol, and the cell eventually regained its polarized morphology and migration behavior. Left panels: Opto-ACTU+, right panels: DIC; untagged cAR1-CIBN was expressed. Top left corners show time in mm:ss format in (A) and in hh:mm:ss in (B).

**Video S11. Cell migration mode and morphology changes in a PTEN– *Dictyostelium* cell, pretreated with PI3K inhibitor LY294002, upon global recruitment of Opto-ACTU+.** Left panels show Opto-ACTU+ and right panels show DIC channel. Untagged cAR1-CIBN was expressed. Top left corners show time in mm:ss format. The 488 nm laser was turned ON to initiate the recruitment, as depicted by the appearance of white text “488 nm ON” in the bottom middle.

**Video S12. Optically confined recruitment of Opto-ACTU– can locally suppress protrusions in RAW 264.7 macrophages.** In both (A)-(C), 488 nm laser was selectively irradiated inside the white rectangular boxes; untagged CIBN-CAAX was expressed; top left corners show time in mm:ss format. Either Opto-ACTU-(A and B) or Opto-CTRL (C) was first locally recruited and then cells were globally stimulated with C5a receptor agonist. The appearance of white-colored text “+C5aR Agonist” in the videos denoting the addition of the agonist. Left panels: Lifeact-mVenus; Right panels: Opto-ACTU– (A and B) or Opto-CTRL (C).

**Video S13. Global recruitment of Opto-ACTU– to the membrane in MCF 10A epithelial cells can deactivate the EGF induced ERK activation.** (A) Translocation of ERK-KTR from cytosol to nucleus upon Opto-ACTU-recruitment. Left panel: ERK-KTR-iRFP713, right panel: Opto-ACTU–. (B) ERKKTR maintained its cytosolic distribution upon Opto-CTRL recruitment. Left panel: ERK-KTR-iRFP713, right panel: Opto-CTRL. In both (A and B), top left corners showing time in hh:mm:ss format. 488 nm laser was globally turned ON at t=0 (as shown by the appearance of green-colored text “488 nm ON”). Cells were pre-treated with a saturated dose of EGF and they were maintained in that condition throughout the experiment.

## METHODS

### Cell culture

The wild-type *Dictyostelium discoideum* cells of axenic strain AX2 were cultured in HL-5 media at 22 °C. Hygromycin (50 μg/mL) and/or G418 (30 μg/mL) were added to the media to maintain cell lines expressing different constructs. PI3K 1^-^/2^-^ and PTEN– *Dictyostelium* cells were cultured like AX2 whereas heat-killed *Klebsiella aerogenes* were added in the culture media to grow PKBA^-^/ PKBR1^-^ *Dictyostelium* cells. Cells were usually maintained in petri dishes and were transferred to shaking culture around 2-4 days before electrofusion or differentiation experiments. PTEN– *Dictyostelium* cells were always grown in petri dishes. All the experiments were done within 2 months of thawing the cells from the frozen stocks.

RAW 264.7 macrophage-like cells were obtained from N. Gautam laboratory (Washington University School of Medicine in St. Louis, MO) and mammary epithelial MCF-10A cells were obtained from M. Iijima laboratory (Johns Hopkins University School of Medicine, MD). RAW 264.7 cells were grown in Dulbecco’s modified Eagle’s medium (DMEM) containing 4500mg/L glucose, L-glutamine, sodium pyruvate, and sodium bicarbonate (Sigma-Aldrich; D6429), supplemented with 10% heat-inactivated fetal bovine serum (ThermoFisher Scientific; 16140071) and 1% penicillin-streptomycin (ThermoFisher Scientific; 15140122). MCF-10A cells were cultured in DMEM/F-12 medium with GlutaMAX (ThermoFisher Scientific;10565042) supplemented with 5% heat-inactivated horse serum (ThermoFisher Scientific; 26050088), 1% penicillin-streptomycin, epidermal growth factor (EGF) (20 ng/ml) (Sigma-Aldrich; E9644), cholera toxin (100 ng/ml) (Sigma-Aldrich; C8052), hydrocortisone (0.5 mg/ml) (Sigma-Aldrich; H0888), and insulin (10 μg/ml) (Sigma-Aldrich; I1882). All cells were subcultured every 2-6 days using cell lineage-specific techniques to maintain healthy confluency. All experiments were done with low passage number cells. All mammalian cells were maintained under humidified conditions at 37 °C and 5% CO_2_.

### DNA constructs

All sensors and actuators were codon-optimized if used for heterologous expression. R(+8)-Pre was obtained from the C-terminal tail of KRas4b and all the serines and threonines were mutated to alanine to prevent phosphorylation and all the lysines were mutated to arginines to avoid ubiquitination^22^. In R(+7)-Pre, R(+4)-Pre, and R(+2)-Pre, arginines are sequentially mutated to glutamines to reduce the charge. The cAR1 is a *Dictyostelium* protein that works as a GPCR. Palm/Pre is the C-terminal tail of HRas. PM-LYN is the first 11 amino acids of the human Tyrosine-protein kinase LYN. In PTEN_1-18_-CAAX, a CAAX motif was added to the first 18 amino acids PTEN. RacG_CT_ is the C-terminal tail of RacG which we identified when we blasted the KRas4b tail in NCBI Protein BLAST (blastp), with organism specified. Optogenetic actuator Opto-ACTU+, which has a net charge +16, was designed by removing CAAX tail from R(+8)-Pre (so that it becomes cytosolic) and making a dimer of it, joined by a GGG linker. Optogenetic actuator Opto-ACTU**–**, which has a net charge −14, was designed using the C-terminal polyanionic tail sequence of mouse protein Rad17.

All surface charge sensors and actuators, i.e. GFP-R(+8)-Pre, GFP-R(+7)-Pre, GFP-R(+4)-Pre, GFP-R(+2)-Pre, GFP-Palm/Pre, GFP-PTEN_1-18_-CAAX, GFP-RacG_CT_, Opto-ACTU+, and Opto-ACTU-were generated by annealing the appropriate synthetic oligonucleotides, followed by restriction enzyme mediated digestion and subcloning into proper *Dictyostelium* or mammalian vectors. All other constructs were made by PCR amplification followed by standard restriction enzyme cloning or by site-directed mutagenesis kit (QuickChange II, Agilent Technologies, 200523). All oligonucleotides were obtained from Sigma-Aldrich. All sequences were verified by diagnostic restriction digest and by standard Sanger sequencing (JHMI Synthesis & Sequencing Facility).

The following plasmid constructs were made in this study. Selected will be deposited on Addgene/dictyBase and rest will be available from the authors upon direct request: a) GFP-R(+8)-Pre (pDM358), b) GFP-R(+8)-Pre (pDEXG), c) GFP-R(+7)-Pre (pDEXG), d) GFP-R(+4)-Pre (pDEXG), e) GFP-R(+2)-Pre (pDEXG), f) GFP-PTEN_1-18_-CAAX (pDEXG), g) GFP-RacG_CT_ (pDEXG), h) GFP-LactC2 (pTX-GFP), i) GFP-Spo20 (pDEXG), j) Opto-ACTU+ (pCV5), k) Opto-CTRL (pCV5), l) cAR1-CIBN (pDM358), m) N150_PKBR1_-CIBN (pDM358), n) Opto-ACTU**–** (pmCherryN1), o) PH_Crac_-Halo (pCV5), p) LimE_Δcoil_-Halo (pCV5).

GFP-R(+8)-Pre (mammalian) was from S. Grinstein (Addgene plasmid # 17274), GFP-LactC2 (mammalian) was from S. Grinstein (Addgene plasmid # 22852), GFP-Spo20, originally made by Vitale et al. ^50^, was a kind gift from G. Du (McGovern Medical School, UTHealth), mPlum-LimE_Δcoil_ was from A. Müller-Taubenberger (LMU Munich), pMD2.G was from D. Trono (Addgene plasmid # 12259), pMDLg/pRRE was from D. Trono (Addgene plasmid # 12251), pRSV-Rev was from D. Trono (Addgene plasmid # 12253), pLJM1-EGFP was from D. Sabatini (Addgene plasmid # 19319), pCRY2PHR-mCherryN1 (i.e. mammalian Opto-CTRL) was from C. Tucker (Addgene plasmid # 26866), CIBN-CAAX was from P. De Camilli and O. Idevall-Hagren (Addgene plasmid # 79574), Lifeact-mVenus was from J. Zhang (Addgene plasmid # 87613), and ERKKTR-iRFP713 was from J. Toettcher (Addgene plasmid # 111510). Rest of the plasmids used in this study were available in Devreotes Lab.

### Transfection

*Dictyostelium* AX2 cells were transfected by standard electroporation protocol. Briefly, for each transfection, 10^7^ cells were pelleted, washed twice with ice-cold H-50 buffer, and subsequently resuspended in 100 μL ice-cold H-50 buffer. Around 1-5 μg of total DNA was mixed with the cell suspension and it was transferred to an ice-cold 0.1 cm gap cuvette (Bio-Rad; 1652089) for electroporation at 850V and 25 μF twice, in a 5 second interval (Bio-Rad Gene Pulser Xcell Electroporation Systems). After a 5 min incubation on ice, electroporated cells were transferred to a 10 cm Petri dish containing HL-5 media supplemented with heat-killed *Klebsiella aerogenes* bacteria. Cells were selected by adding hygromycin B (50 μg/ml) and/or G418 (20-30 μg/ml) after 1-2 days, as per the antibiotic resistances of the vectors. For optogenetics and chemically-induced-dimerization experiments where three different protein co-expressions were necessary, two different pCV5 vectors and one pDM358 vector were used and cells were selected with both drugs.

RAW 264.7 cells were transfected by nucleofection in an Amexa Nucleofector II device, using Amexa Cell line kit V (Lonza; VACA-1003), following a pre-existing protocol^101^. For each transfection, 3×10^6^ cells were harvested, resuspended in 100 μL supplemented Nucleofector Solution V. Then total 4-6 μg of DNA mixtures were added and immediately transferred to a Lonza cuvette for electroporation using program setting D-032. 500 μL of pre-warmed pH adjusted culture media was added to electroporated cells in the cuvette subsequently. Cell suspension was then transferred to a 1.5 mL vial and incubated at 37 °C and 5% CO_2_ for 10 mins. Next, cells were allowed to attach on coverslip chambers for an hour. 2 mL of pre-warmed pH adjusted culture media was added to each chamber and cells were further incubated for 4-6 hours before imaging.

MCF-10A cells were transiently transfected using Lipofectamine 3000 Transfection Reagent (ThermoFisher Scientific; L3000001), as per manufacturer’s protocol. Briefly, ∼0.8 µg DNA was mixed in 160 µL serum-free Opti-MEM (ThermoFisher Scientific; 31985062) media containing 3 µL Lipofectamine 3000 reagent and incubated for 5-7 mins at RT to allow formation of DNA-lipid complex. Post-incubation, DNA-lipid complex was added to 2×10^5^ MCF-10A cells plated on a 2-well glass chamber. Cells were incubated at 37 °C/5% CO_2_ for 5 hours, after which the DNA-lipid complex was removed, cells were washed thoroughly, and incubated for a further 16 hours before imaging.

### Drugs and reagents

Annexin V, Alexa Fluor 488 conjugate was obtained from ThermoFisher Scientific (Invitrogen; A13201) and was stored in 4 °C. Latrunculin A (Enzo Life Sciences; BML-T119-0100) was dissolved in DMSO to make a stock solution of 5 mM. Caffeine (Sigma-Aldrich; C0750) was dissolved in ddH_2_O to make a stock solution of 80 mM. Rapamycin (Sigma-Aldrich; 553210) was dissolved in DMSO to prepare a 10 mM stock solution. cAMP (Sigma-Aldrich; A6885) was dissolved in ddH_2_O to make a stock solution of 10 mM. Janelia Fluor 646 HaloTag (Promega Corporation; GA1120) was dissolved in DMSO to prepare 200 μM stock solution which was stored in 4 °C and was diluted 1000X during the experiment. C5a receptor agonist FKP-(D-Cha)-Cha-r (Anaspec; 65121) was dissolved in 1X PBS to make to 2.5 mM stock solution. Anti-BSA antibody was obtained from Sigma (Sigma-Aldrich; SAB4200688). Stock solution for EGF (Sigma-Aldrich, E9644) was prepared by dissolving it in 10 mM acetic acid to a final concentration of 1 mg/ml. Unless otherwise mentioned, everything was stored as small aliquots in −20 °C.

### Microscopy and live cell imaging

Unless otherwise mentioned, all experiments were performed inside a heated (37 °C) chamber with a 5% CO_2_ supply (in case of mammalian cell imaging), or on a 22 °C stage (in case of *Dictyostelium* imaging). All time-lapse live-cell images were acquired using one of these four microscopes: a) Zeiss LSM780-FCS Single-point, laser scanning confocal microscope, (Zeiss Axio Observer with 780-Quasar; 34-channel spectral, high-sensitivity gallium arsenide phosphide detectors), b) Zeiss LSM880-Airyscan FAST Super-Resolution Single-point confocal microscope (Zeiss AxioObserver with 880-Quasar (34-channel spectral, high-sensitivity gallium-arsenide phosphide detectors), c) Zeiss LSM800 GaAsP Single-point, laser scanning confocal microscope with wide-field camera, and d) Nikon Eclipse Ti-E dSTROM Total Internal Reflection Fluorescence (TIRF) Microscope (Photometrics Evolve EMCCD camera).

In Zeiss 780 and 800, 488 nm (argon laser) excitation was used for GFP; 561 nm (solid-state) excitation was used for RFP, mCherry, and mPlum; and 633 nm (diode laser) excitation was used for iRFP713 and Janelia Fluor 646 HaloTag. In Zeiss 880, 488 nm (argon laser) excitation was used for GFP; 514 nm (argon laser) excitation was used for YFP and mVenus; 594 nm (HeNe laser) excitation was used for mCherry; and 633 nm (diode laser) excitation was used for iRFP713. In Nikon TIRF, 488nm (argon laser) excitation was used for GFP and 561 nm (0.5W fiber laser) excitation was used for mCherry and RFP. In Zeiss 780, 800, and 880, 40X/1.30 Plan-Neofluar oil objective (with appropriate digital zoom) was used and in Nikon TIRF, 100x/1.4 Plan-Apo oil objective was used. Both 780 and 880 confocal microscopes are operated by ZEN Black software, 800 confocal microscopes are operated by ZEN Blue software, whereas the TIRF is controlled by Nikon NIS-Elements. To visualize basal/ventral waves in *Dictyostelium* and RAW 264.7 cells, either TIRF microscope was used or confocal microscopes were focused on the very bottom of the cell to capture the substrate-attached surface of the cell.

### Cell differentiation

For *Dictyostelium* cell development, 8×10^7^ cells of exponential growth phase were collected from suspension culture and pelleted. After washing twice with DB (Development buffer; 5 mM Na_2_HPO_4_, 5 mM KH_2_PO_4_, supplemented with 2 mM MgSO_4_ and 0.2 mM CaCl_2_), cells were resuspended in 4 mL DB and shook at 110 rpm for 1 hour. After 1 hour, the shaking was continued, and the cells were pulsed with 50-100 nM of cAMP (5 sec pulse every 6 mins) using a time-controlled peristaltic pump for next 5-6 hours. This allowed the cells to become developed and polarized. After development, from the shaker, around 2-5×10^4^ cells were transferred to an 8-well coverslip chamber, resuspended thoroughly in 450 μL of DB, and incubated for 20-30 min before starting the image acquisition.

### Frustrated phagocytosis and osmotic shock

To visualize ventral waves in RAW 264.7 macrophages, we have slightly modified a pre-existing protocol^44, 102^. Briefly, Nunc Lab-Tek 8-well coverslip chambers were prewashed with 30% nitric acid, coated with 1 mg/mL BSA for 3 hours, washed with PBS, and finally incubated with 5 μg/mL anti-BSA antibody (1:200 dilution) for 2 hours. Chambers were finally washed two times with PBS to remove excess antibodies. Before the imaging, transfected RAW cells were starved in suspension in 1X Ringer’s buffer (150 mM NaCl, 5 mM KCl, 1 mM CaCl_2_, 1 mM MgCl_2_, 20 mM HEPES and 2 g/L glucose, pH 7.4) for 30 min. Next, these cells were introduced to the opsonized chambers and allowed to spread on antibody-coated surface for 5-10 min. To initiate more waves during imaging, hypotonic shock was applied to cells using 0.5X Ringers solution.

### Electrofusion

Exponential growth phase *Dictyostelium* cells from suspension culture were collected, washed, and resuspended in 10 mL SB (17 mM Soerensen buffer containing 15 mM KH_2_PO_4_ and 2 mM Na_2_HPO_4_, pH 6.0) at a density of 1.5 × 10^7^ cells/ml. The cells were rolled gently in a conical tube for around 30-40 min to promote visible cluster formation. 800 μL of rolled cells were transferred to a 4-mm-gap Bio-Rad electroporation cuvette, using pipette tips with cut off edges (to ensure clusters remain intact). The electroporation was performed with 1kV, 3 μF once, then with 1kV, 1 μF twice more (with 3 s gap between two pulses) to facilitate membrane hemifusion. 35 μL of cells were transferred from the cuvette to a Nunc Lab-Tek 8 well chamber and incubated for 5 more min and then fresh SB buffer supplemented with 2 mM CaCl_2_ and 2 mM MgCl_2,_ was gently added to these cells. Cells were incubated at 22 °C for next 1-1.5 hours for recovery before imaging.

### Annexin V binding assay

Growth phase *Dictyostelium* cells were transferred to an 8-well Nunc Lab-Tek coverslip chamber and allowed to adhere for 10-15 min. Next the HL-5 medium was removed and 450 μL DB buffer was added to the cells. Cells were incubated at 22 °C for ∼ 60-120 min. After that, the DB buffer was exchanged with DB buffer with excess calcium (2.5 mM of CaCl_2_ final concentration) and incubated for 30 min. The coverslip chambers were transferred to ice bath (to slow down the protrusion formation and withdrawal frequency) and rest of the procedure was done in ice. Next, 10 μL of Annexin V, Alexa Fluor 488 conjugate was added to each well and it was allowed to bind for 45-60 sec. Then the media (containing Annexin V, Alexa Fluor 488 conjugate) was aspirated and was quickly washed for two times using DB with excess calcium buffer to get rid of all unbound Annexin V. Cells were then fixed without permeabilization, using 2% paraformaldehyde and 0.25% Glutaraldehyde in HL-5 (with 2.5 mM of Ca^2+^) for 15 min and then washed twice and finally put under DB with excess calcium for imaging.

### Gradient stimulation assay

Gradient stimulation assay by cAMP-filled needle was performed as per the established protocol^103, 104^. *Dictyostelium* cells co-expressing PH_crac_-mCherry and GFP-R(+8)-Pre were differentiated (as per above mentioned protocol) for 5.5-6.5 hours and were put under cAMP gradient. Cells were pre-treated with LatA to inhibit cytoskeletal activities. A 10 μM cAMP-filled micropipette connected to a micromanipulator was used to provide the gradient stimulation. A microinjector connected to the micromanipulator was employed in a continuous injection mode with a compensation pressure of 70 hPa.

### Chemically induced dimerization

The plasmids and experimental details of CID system were mostly described in our previous reports^9, 57^. Here, to generate Fig. 2, E to I and fig. S5, E to G, two following combinations were used to express the systems: a) cAR1-FKBP-FKBP (pCV5) as membrane anchor, mCherry-FRB-Inp54p (pCV5) as recruitee, GFP-R(+8)-Pre (pDM 358) as readout; and b) cAR1-FKBP-FKBP (pDM 358) as membrane anchor, mCherry-FRB-Inp54p (pCV5) as recruitee, PH_PLCδ_-YFP (pCV5) as readout. Growth phase *Dictyostelium* cells were transferred to an 8-well coverslip chamber and incubated for 10-15 min so that they can adhere well. Then, the HL-5 medium was replaced with 450 μL of DB buffer. After 15-20 min of media change, image acquisition was started. To felicitate recruitment of Inp54p to plasma membrane, after imaging a certain number of frames, rapamycin was added gently to the chamber (to final concentration 5μM) during image acquisition. In case of Fig 2I, first it was ensured that the Inp54p was recruited to the membrane by looking at a few confocal slices at the middle of the cell, then focused to the substrate-attached surface to visualize waves.

### Optogenetic regulation of cell migration

To analyze the effect of surface charge lowering on cell polarity and migration pattern, *Dictyostelium* cells was selected against both Hygromycin and G418 to co-express LimE_Δcoil_-Halo (pCV5), cAR1-CIBN (pDM 358), and Opto-ACTU+ (pCV5) or Opto-CTRL (pCV5). Cells were properly developed as described in “Cell differentiation” section. After 6-7 hours of development, around 2-5×10^4^ cells were collected from the shaker and transferred to an 8-well coverslip chamber. Then cells were resuspended thoroughly in 450 μL of DB, and incubated for around 20 min before starting the image acquisition. For experiments presented in Figures 5G, 5H and Figures S10E-S10L, developed *Dictyostelium* cells were incubated with 40 μM LY294002 or 20 μM PP242 or both for >45 min in DB buffer before image acquisition was started. For global recruitment experiments, after imaging ≥320 s, the 488 nm Laser was turned ON globally to initiate recruitment and was intermittently turned on at a lower intensity during image acquisition (usually for around 970 ms after each 8s) to keep Opto-ACTU+ or Opto-CTRL on the membrane throughout the imaging. For spatially confined optical recruitment, a region of interest was drawn and that particular area was illuminated was 488nm laser in multiple iteration. To analyze the effect of surface charge elevation on cell polarity and migration pattern, RAW 264.7 macrophage cells co-expressing Lifeact-mVenus, CIBN-CAAX, and Opto-ACTU**–** or Opto-CTRL was used. After 4-6 hours of nucleofection (as described in “Transfection” section), media was aspirated from chambers and cells were put in 450 μL of new warm HBSS buffer (Hank’s Balanced Salt Solution) containing 1 g/L of glucose and cells were incubated for another 0.5-1 hour before starting image acquisition. During imaging, first Opto-ACTU**–** or Opto-CTRL was selectively recruited first using 488 nm laser (as described above) and subsequently C5a receptor agonist FKP-(D-Cha)-Cha-r was added (diluted in HBBS; final concentration of 10 μM) and image acquisition was continued.

### Optogenetic deactivation of ERK

The KTR technology was developed to convert kinase activities into a nucleocytoplasmic shuttling equilibrium of a genetically coded sensor so that the kinase activity can be visualized real time and quantitated^105, 106^. Basically, ERKKTR sensor becomes cytosolic to nuclear upon the deactivation of ERK. To analyze the effect of optogenetic perturbation of negative surface charge of the inner membrane, MCF10A cells were transfected with ERKKTR-iRFP713 (as readout), CIBN-CAAX (as membrane anchor), along with Opto-ACTU**–** or Opto-CTRL (as recruitee). Transfected cells were incubated overnight in complete culture medium containing 5% horse serum,10 μg/ml, and 20 ng/mL EGF, which facilitated activation of ERK. Imaging was performed in the same media. After 25 mins of image acquisition, 488 nm Laser was globally turned ON to facilitate recruitment and was intermittently turned on during image acquisition to keep Opto-ACTU**–** or Opto-CTRL on the membrane throughout the imaging. ERKKTR localization profile was used to quantitate ERK activity over the time.

### Image analysis

Image analysis was performed in MATLAB 2019a (MathWorks, Natick, MA, USA) and Fiji/ImageJ 1.52i (NIH). The results were plotted using MATLAB 2019a, OriginPro 9.0 (OriginLab, Northampton, MA, USA), or GraphPad Prism 8 (GraphPad Software, San Diego, CA, USA).

### Colocalization study

Image analysis for the colocalization study was performed with custom codes written in MATLAB 2019a (MathWorks, Natick, MA, USA). As the preprocessing steps, the background subtraction and the gaussian/top hat filtering were applied to all the images. The cell area from the background and the bright patches of the protein of interest from the cell area were segmented using thresholding method.

To quantify the extent of colocalization between two proteins of interest, say A and B in a cell, the following conditional probabilities were computed:

1. Prob(A_high_|B_high_): the probability of finding ‘high localization’ of protein A in the regions of ‘high localization’ of protein B.
2. Prob(B_high_|A_high_): the probability of finding ‘high localization’ of protein B in the regions of ‘high localization’ of protein A.
3. Prob(A_high_|B_low_): the probability of finding ‘high localization’ of protein A in the regions of ‘low localization’ of protein B.
4. Prob(A_low_|B_high_): the probability of finding ‘low localization’ of protein A in the regions of ‘high localization’ of protein B.
5. Prob(B_high_|A_low_): the probability of finding ‘high localization’ of protein B in the regions of ‘low localization’ of protein A.
6. Prob(B_low_|A_high_): the probability of finding ‘low localization’ of protein B in the regions of ‘high localization’ of protein A.

High and the low localization of a protein is decided by the threshold values assumed in the segmentation step. If both the proteins, A and B were colocalized in a cell, then the following inequalities would hold true:

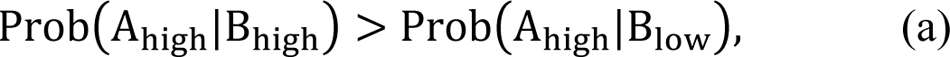

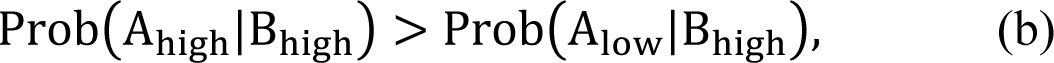

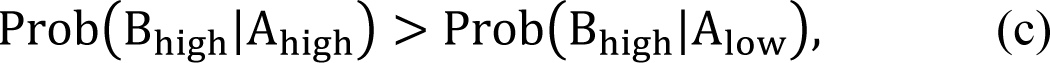

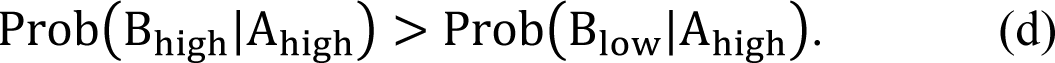

The inequality sign would be preserved under the change of ordering. The sign reverses if the proteins do not colocalize. Clearly in case of the colocalization, the ratio of these conditional probabilities (from Eqn. (a-d)) will be greater than 1, whereas it will be less than 1 for the cases of complementary localization. Taking logarithm of these ratios further differentiates these two cases leading to positive and negative values, respectively. For the concise representation of the data, we computed the following average of the four ratios of conditional probabilities for each image frame as follows, and named it as Conditional Probability Index (henceforth CP index):

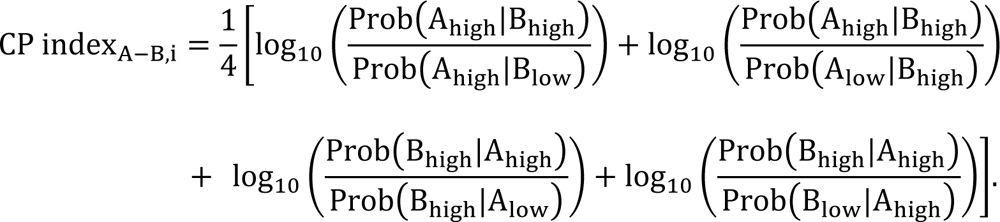

For the box and whisker plot representation, we averaged the CP index. over *n* frames analyzed for every cell image (henceforth Avg. CP index.) as follows:

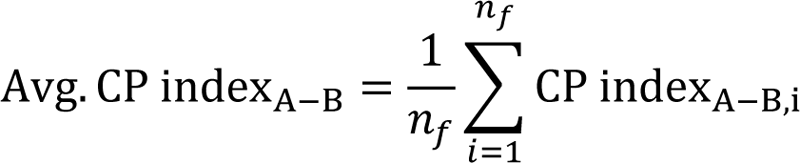

where *n_f_* (≥ 20) is the total number of frames analyzed for a sample.

### Kymographs

For the membrane kymographs the cells were segmented against the background following standard image processing steps with the custom code written in MATLAB 2019b. The kymographs were created from the segmented cells as previously described (10). A linear colormap was used for the normalized intensities in the kymographs. For grayscale kymographs, black indicating the lowest intensity and white the highest whereas in colored kymographs, lowest intensity was indicated by blue and highest by yellow.

Line kymographs that accompanied ventral waves were generated in Fiji/ImageJ by drawing a thick line with line width 8-12 pixel and processing the entire stack in “KymographBuilder” plugin.

### Line scan intensity profile

Line scans were performed in Fiji/ImageJ (NIH) by drawing a straight-line segment (using line tool) inside the cells with line width 8-12 pixel so that we can obtain an average intensity value and one particularly bright or dim pixel does not skew the result. The intensity values along that particular line was obtained in green as well as red channels using “Plot Profile” option. The values were saved and normalized, in OriginPro 9.0 (OriginLab, Northampton, MA, USA). Then the intensity profiles were graphed and finally smoothened using Savitzky-Golay or Adjacent-Averaging method, using proper boundary conditions. For a particular linescan, the green and red profiles were smoothed using exact same parameters to maintain consistency.

### Time-series plot of Membrane/Cytosol ratio

The cells were segmented into membrane and cytosolic masks following standard image processing steps with the custom code written in MATLAB 2019b. The average intensities from the channels were computed for both the masks. The computed intensities were later corrected for the photobleaching effect by dividing the values by an exponential fit to the temporal profiles of the respective normalized average intensities in the cell. Finally, we computed the ratios of the corrected average intensities for different channels. For the plotting of the temporal profiles of the red channel data we normalized the pre-recruitment average profiles to unity, whereas for the green channel, the normalization was done using the respective steady-state value.

### Cell tracking and migration analysis

To segment the cells, first, using the “Threshold” option of Fiji/ImageJ, image stack was threholded based on a proper threshold value so that the generated binary image covers all the pixels of the cells. Range was not reset and “Calculate threshold for each image” option was unchecked. Subsequently, using “Analyzed Particles” option, a size-based thresholding was applied and cell masks were generated so that small non-cell particles could be excluded from further analysis. Next, “Fill holes”, “Erode”, and “Dilate” options were applied sequentially (for a couple of times, in some cases) to obtain the proper binarized mask for cells. To generate the migrating cell outlines in Fig. 3I, “Outline” command was operated on binarized masked cells and finally “Temporal-Color Code” was used. For other plots, “Shape descriptors” and “Centroid” options was checked inside “Set measurements” menu and “Analyzed Particles” option was used, this time to obtain the values of centroid coordinates and circularity values (circularity is 4*π*area/perimeter^2^). Then from the replicates of circularity values, mean and SEM were determined in GraphPad Prism and they were exported to OriginPro for plotting. For the centroid values, the starting point was set to origin for each track. Basically, *x*^′^_i_ = (*x*_i_ − *x*_0_) and *y_i_^′^ = (y_i_− y_0_)* were applied for *i=0 to (n-1).* The new translated coordinates were plotted in OriginPro to generate the tracks. Same increment colormap was used in before-recruitment and after-recruitment tracks so that the tracks can be visually compared in a pairwise fashion in Fig. 3F, fig. S7E, and fig. S9B. To get the velocity values, displacement between each two frames were computed using 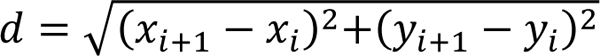 formula. This displacement was divided to time interval to compute speed for each case and these speeds were time-averaged over the frames to generate each datapoint for the migration speed box plots. Based on those data points, in GraphPad Prism, the box and whiskers were plotted.

### Global protrusion formation analysis

The new protrusion formation frequency in *Dictyostelium* cells upon global recruitment of Opto-ACTU+ or Opto-CTRL was computed in Fiji/ImageJ. In the stack of the binarized cell masks (as generated in “*Cell tracking and migration analysis*” section), “Stack Difference” command of “Multi Kymograph” was applied. The stack was inverted using “Invert” command and a size-based thresholding was performed using “Analyze Particles” command. If required, the resulting stack was processed further using “Despeckle” command and then protrusion numbers were manually counted. The numbers were counted for the same frames for which the migration speed was quantified. To calculate the fraction of membrane state showing front activity in HL-60 neutrophils, we first segmented the cells and binarized cell masks were generated by sequentially applying “Threshold”, “Analyze Particles”, “Fill holes”, “Erode”, and “Dilate” options (as described above in “*Cell tracking and migration analysis*” section). The “Perimeter” option was checked in “Set Measurements” menu and the perimeter of all the cell images the stacks were obtained by applying “Analyze Particles” command on binarized cell masks. To get the front or protrusion area, first Lifeact channel was binarized again with a different lower threshold value (usually 1.5-2 times of the value that was used to binarize the whole cell). On the new stack with masks, “Erode” and “Dilate” command was applied and a size-based thresholding was performed using “Analyze Particles” command. The “Skeletonize” command was applied to obtain single-pixel-wide shapes and the skeletonize stacks were converted to a time-series. A custom-written Python script was run using the Fiji/ImageJ Jython interpreter to process the time-series stack so that we can obtain the “longest shortest paths” using the “Analyze Skeleton” option. Finally, using a custom-written MATLAB code, the longest shortest paths were added for each frame. This was divided by the perimeter values to determine the fraction of front in each frame and then a time-average was determined for required time intervals to generate each data point for the box and whisker plot. All box plots were generated in GraphPad Prism.

### Selective protrusion formation analysis

To perform the selective protrusion formation analysis upon optically confined recruitment of Opto-ACTU+, Opto-ACTU**–**, and Opto-CTRL, we segmented the recruitment area on the membrane with proper threshold. We then marked the area using the segmented line tool. Using a custom-written ImageJ Macro, “Fit spline” and “Straighten” commands were sequentially operated. The macro was then used to find and mark the midpoint. The centroid of the entire cell was also found and marked using another macro. Using the angle tool, considering the centroid as vertex, the angles between protrusions and the midpoint of the recruitment areas were obtained. Values were imported to MATLAB and plotted using the “polarhistogram” command. Sturges’ formula was used to determine the minimum number of bins.

### KTR translocation analysis

The nucleus to cytosol ratio of ERKKTR was computed over the time to quantitate the extent of ERK deactivation upon Opto-ACTU**–** or Opto-CTRL recruitment. The cells were first segmented in Fiji/ImageJ by sequentially applying “Threshold”, “Analyze Particles”, “Fill holes”, “Erode”, and “Dilate” options (as described in detail in “*Cell tracking and migration analysis*” section).

Using “Analyze Particles” option, the whole cell intensities were obtained. Nucleus intensities was obtained manually by selecting nucleus from the phase channels of the stack. The cytosolic intensities were found by subtracting nuclear intensity from the whole cell intensity. After ratios were found, a conditional formatting rule was applied in Microsoft Excel to generate heat maps in fig. S12, B and C. The ratios were further imported to GraphPad Prism to calculate mean and SEM and those are plotted in OriginPro to generate the time-series of plot Fig. 4I.

### Computational model

#### Excitable signal transduction system

The excitable system consists of three states – F, B, and R. The system dynamics can be described by the stochastic partial differential equations (SPDE) as follows:

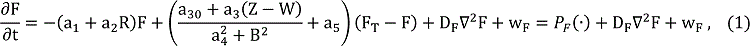

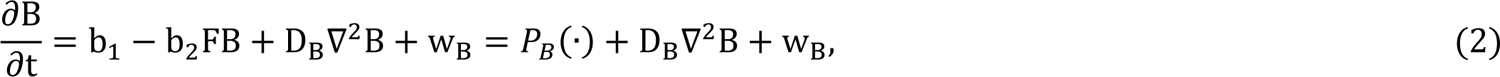

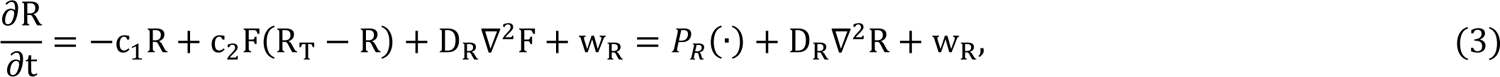

where F_T_, R_T_ are the total amount of F and R, respectively. The stochasticity affecting state X is introduced through the signal, w_X_ representing the white noise process with respective variance σ_X_^2^∝ P_X_(·). Eqn. (1-3) is the manifestation of the standard activator-inhibitor system where F is the fast activator and R is the slow inhibitor. The original auto catalytic loop around the activator is replaced by the mutual inhibitory interactions between F and B. First two reaction terms in Eqn. (1) represent the basal and R-dependent inactivation of F, respectively. The third reaction term represents the nonlinear activation with a negative feedback from B along with the positive feedback from the downstream polarity module (discussed later). The fourth term involves the linear basal activation rate. The reaction terms in Eqn. (2) involve basal activation and F-dependent deactivation of B, respectively. In the Eqn. (3), the reaction terms represent the deactivation and F-dependent activation of R. The total amounts of F and R are conserved in the system.

#### Polarity Mechanism

To simulate the behavior of polar cells, we coupled the excitable system with a two-component biasing network based on the local excitation and global inhibition (LEGI) principle ^84^:

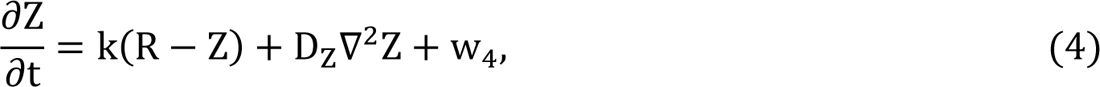

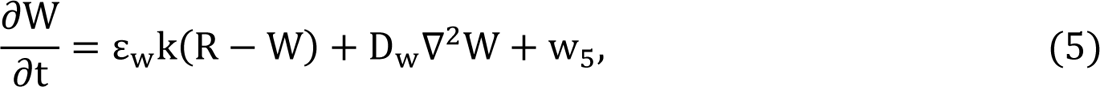

where Z is the fast, local activator and W is the slow (ε_w_ < 1), globally diffusing (D_w_ ≫ D_Z_) inhibitor. Both activator and the inhibitor of the polarity mechanism are activated by R of upstream excitable system. The output of this biasing network (Z − W) positively affects the nonlinear activation of F thus increasing the probability of firing at the former firing locations and resulting long lived patches of F activity.

#### Positively Charged Actuator Dynamics

To model the partitioning of the uniformly recruited charge sensors on the membrane, we considered two populations of charge sensors – fast diffusing, C_mf_ and slow diffusing C_mb_ (D_Cmf_+ ≫ D_Cmb_+). The system dynamics assumed for the positive charge actuator (Opto-ACTU+) is given by,

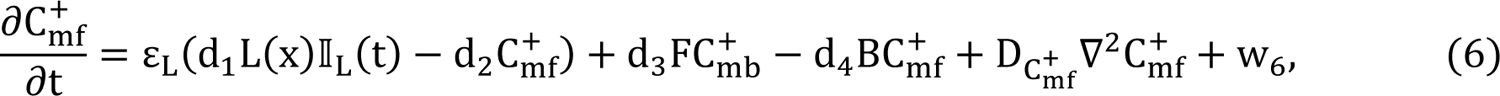

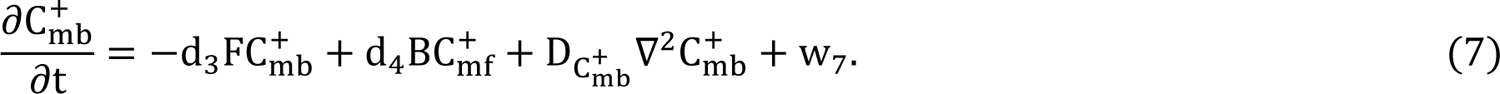

The first term in Eqn. (6) represents the spatiotemporal recruitment of charge actuators to the membrane. The indicator function L(x) controls the spatial location(s) of the recruitment whereas ∥_L_(t) controls the instant and the time span. The second term denotes the reverse process. The third and the fourth term of Eqn. (6) represent F and B-mediated interconversions of *C_mf_*^+^ and C_mb_^+^. The total positive charge actuators (C_m_^+^) on the membrane is given by, C_m_^+^ = C_mf_^+^ + C_mb_^+^. The effect of positive charge actuation on the membrane states is incorporated through a negative feedback term involving C_mb_^+^ in the B dynamics as follows:

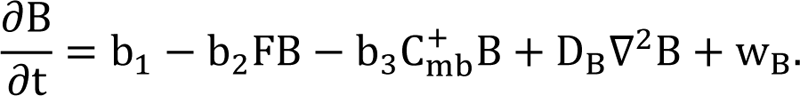

#### Negatively Charged Actuator Dynamics

For modeling negative charge actuators, we followed a similar methodology as discussed above. The system dynamics assumed for the negative charge actuator (Opto-ACTU-) is given by,

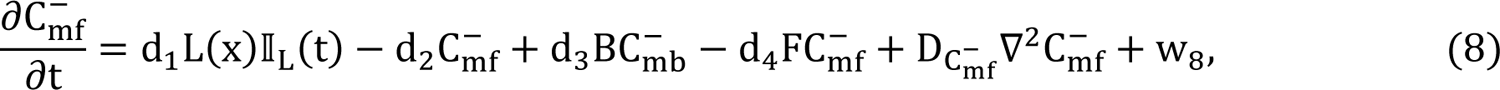

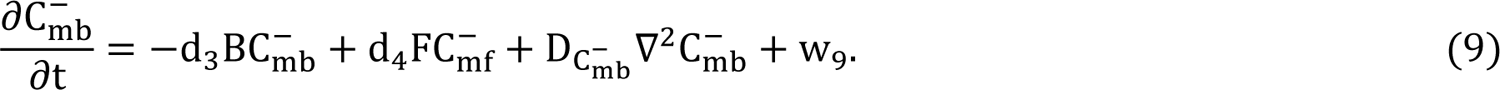

The total negative charge actuators (C_m_^−^) on the membrane is given by,C_m_^−^ = C_mf_^−^ + C_mb_^−^. The effect of negative charge actuation on the membrane states is incorporated through a negative feedback term involving C_mb_^−^ in the F dynamics as follows:

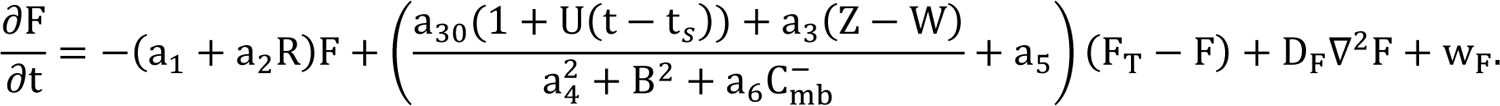

To mimic the experimental conditions, we reduced the contribution of the polarity mechanism and added an additional term, U(t − t_S_) denoting the global agonist stimulation as follows:

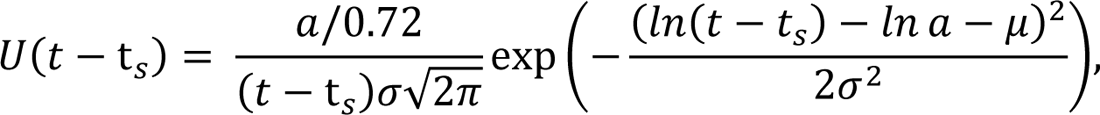

where t_S_ denotes the time instant of application of the global stimulation.

For the simulation a 1D domain of length 30 A.U was chosen with periodic boundary condition. The parameter values (in A.U.) used in these simulations are provided in Table S2. With the central difference approximation of the spatial second derivatives, the coupled SPDEs were converted to a system of coupled stochastic ordinary differential equations (ODE). The stochastic ODEs were numerically solved using Euler-Maruyama method in MATLAB 2019b (MathWorks, Natick, MA, USA) with the aid of the SDE toolbox. The time and space discretization steps were chosen to be Δ*T* = 0.001 A.U. and Δ*x* = 0.1 A.U. respectively. All the states were restricted to be strictly nonnegative.

### Statistical analysis

The statistical analyses were performed using MATLAB 2019a (MathWorks, Natick, MA, USA) and GraphPad Prism 8 (GraphPad Software, San Diego, CA, USA). Time-series data was shown as mean ± SEM or mean ± SD, as indicated in the figure captions. All box plots were graphed following Tukey’s convention. Statistical significance and P-values were determined by paired or unpaired two-tailed non-parametric tests as specifically indicated in the figure captions. The following convention was followed to show P-values: ns denotes P>0.05, * denotes P ≤ 0.05, ** denotes P ≤ 0.01, *** denotes P ≤ 0.001, and **** denotes P ≤ 0.0001.

## Data availability

All data needed to evaluate the conclusions in the paper are present in the main text or the supplementary materials. Any additional requests for information or data will be fulfilled by the corresponding author upon reasonable request.

## Code availability

Computational simulation codes will be available upon reasonable request.

